# The longitudinal single-cell panorama of cynomolgus monkey ovary throughout lifespan revealed a conserved ovarian clock between primates

**DOI:** 10.1101/2023.08.14.553309

**Authors:** Long Yan, Xin Long, Yan Zhao, FeiYan Zhao, Wan Tu, Qiuyun Yang, Jingjing Qian, Jinglei Zhai, Meijiao Wang, Yuqiong Hu, Beijia He, Youqiang Su, Xiangxiang Jiang, Fei Gao, Hongmei Wang, Fan Guo

**Affiliations:** State Key Laboratory of Stem Cell and Reproductive Biology, Institute of Zoology, Chinese Academy of Sciences, Beijing 100101, China; Institute for Stem Cell and Regeneration, Chinese Academy of Sciences, Beijing 100101, China; Beijing Institute for Stem Cell and Regenerative Medicine, Beijing 100101, China; University of Chinese Academy of Sciences, Beijing 100049, China; West China School of Basic Medical Sciences & Forensic Medicine, Sichuan University, Chengdu, Sichuan 610041, China; Shandong Provincial Key Laboratory of Animal Cells and Developmental Biology, School of Life Sciences, Shandong University, Qingdao 266237, China; NHC Key Laboratory of Study on Abnormal Gametes and Reproductive Tract, Anhui Medical University, Hefei 230032, China

**Author notes:** Correspondence: Fei Gao, Hongmei Wang, Fan Guo. These authors contributed equally to this work (Long Yan, Xin Long, Yan Zhao).

## Abstract

Ovarian function is critical for female fertility and impacts reproductive longevity. It is of great importance to accurately predict the aging process within the ovary for fertility assessment and disease diagnosis. However, cell metrics for evaluating the ovarian aging rate are still in urgent need, and molecular insights into ovarian development and dysfunction during the primate life course are also limited. Here, we reported large-scale ovarian cell atlas of consecutive development of cynomolgus monkeys across 22 years with 20 time points, covering the foetal, newborn, prepubertal, pubertal, adult, perimenopausal and menopausal stages. We characterized and validated distinct molecular signatures of each cluster of cells within primate ovaries, and uncovered a previously undocumented RHOXF1-positive oocyte type during primordial follicle assembly in primates. Furthermore, the constitution and developmental trajectories of primate germ cells, granulosa cells and stromal / theca cells were also elucidated, and their precursors were identified. More importantly, dynamics of cellular compositions were unravelled through the ovarian development, featured by granulosa, epithelial, stromal, and immune cells that showed strong temporal heterogeneity spanning lifetime, whilst referred to the key function during the corresponding stages. Based on the correlations of each cell type with age and stage-specific molecular dynamics, we further constructed a transcriptomic ovarian clock which could perceive an effective biological age prediction of the ovary and further applied to humans. The findings reveal granulosa, epithelial, and stromal cells as the highest performance predictors of ovarian biological age, while highlighting the crucial role of AGE-RAGE and Relaxin signaling pathways in regulating ovarian aging. Our work not only provide valuable resource for obtaining insights into the development, aging and dysfunction of key organs, but also establish a transcriptomic clock to predict biological ovarian aging thus to be potential clinical implementation in future.

## Introduction

To obtain the insights of aging process and the adverse effects of reproductive senescence, it is indispensable to map spatio-temporal development of specific organ to clarifying the biological changes and molecular parameters over time. To date, human organ development throughout the course of life is largely constrained due to the lack of access to intact and healthy tissues. Nonhuman primates (NHPs) are believed to be optimal surrogates (*1–3*); however, few studies have been reported, and these reports had finite developmental time points and small sample sizes (*4–6*). The ovary is crucial for female fertility and health (*7*), the source organ for oogenesis (*8*) and an endocrine tissue that secretes hormones to affect the whole body (*9*). Knowledge of cellular diversity and its developmental trajectory is pivotal for understanding ovarian function and female fertility. In addition, the maturation of oocytes involves the orchestration of multiple cell types within the ovary and the somatic-cell microenvironment (*10, 11*), while insights into cell-cell interactions among ovarian cells are lacking. Notably, understanding the establishment of ovarian follicle pools, which impacts a female’s fertility for life time, requires the decipherment of molecular profiles of oocytes and their interactions with surrounding granulosa cells (GCs) during primordial follicle (PF) assembly and growth. Although the constitution and molecular profiles of ovarian cells in different stages of development have been extensively studied in mice (*12–15*), these parameters are poorly delineated in primates.

There is a growing interest to develop reliable and generalize tools for aging prediction and disease diagnosis in recent years. Biological aging rates are highly variable among individuals of the same chronological age, it is crucial to assess the biological rather than chronological age when quantify aging, especially for reproductive organs. Advanced studies had present deep learning models that successfully predict senescence or diagnose diseases based on image and video data (*16, 17*). However, the vast majority of the studies based on DNA methylation (called epigenetic clocks) meta-analysis (*18–20*), and age-related methylation changes do not overlap with transcriptomic changes in human bulk tissue (*21, 22*). The general lack of understanding of the molecular and cellular causes and consequences of genomic CpG methylation remains a barrier to realizing the potential of these models (*23*). A single-cell transcriptomic atlas across the lifespan would lend greater insights than a static genetic investigation and would benefit the study of biological aging and characterize somatic cells and universal markers suitable for predicting senescence and even risk of disease (*24, 25*). Although single-cell transcriptome analysis of ovarian tissues from adult humans has been reported, these tissues were mainly derived from cancer patients and were subjected to cryopreservation or long-term androgen treatment before characterization (*26, 27*); thus, the influences of diseases or chemicals could not be excluded. Recently, a report of single-cell transcriptomes of 2,183 ovarian somatic cells from two time points (young and aged cynomolgus monkeys) examined alterations mainly in GCs between the aging and control groups (*28*); however, many functional somatic cell types, such as endocrine theca cells, were not detected in this dataset (*28*). Additionally, the limitations in cell numbers prevented a detailed analysis in subgroups of ovarian somatic cells (*28*), so the similarities of ovarian cells between humans and NHPs could not be systematically examined, leaving the question of how cell constitutions and molecular markers are conserved between these species. Notably, the progenitor cells and their developmental trajectory in the primate ovary still need to be identified, which warrants the study of ovary development and dysfunction throughout the life cycle.

In this study, we generated a comprehensive single-cell atlas of cynomolgus monkey ovaries from the embryonic stage to natural menopause, covering seven critical periods across 22 years, and investigated the molecular and cellular processes underlying the developmental path, aging and diseases. By applying integrated analysis to the distinct molecular signature of different ovarian cell types throughout the whole lifespan of cynomolgus monkey, we successfully developed high-reliability transcriptomic clocks which were conserved among primates for predicting the pace of ovarian biological aging.

## Results

### Single-cell atlas of the ovaries throughout the fertility cycle of the cynomolgus monkey

To investigate the dynamics of the constitution of ovarian cells and the cell-cell interactions during primate development, we collected ovaries from 17 cynomolgus monkeys and performed single-cell RNA sequencing, and integrated two published datasets including five foetal ovaries from ourselves (*29*) and the other study (*30*) (Fig. 1A). The ovarian samples used in this study covered key stages of development and fertility periods of the cynomolgus monkey, including the foetal period, newborn period, prepuberty, puberty, adult period, perimenopause and menopause (*31, 32*). All ovaries were confirmed by their morphologies and exhibited age-related follicle reserve dynamics (Fig. 1B and fig. S1A). Furthermore, ovarian fibrosis increased with age, as reported previously (*33, 34*) (fig. S1, B and C).

**Fig. 1.**
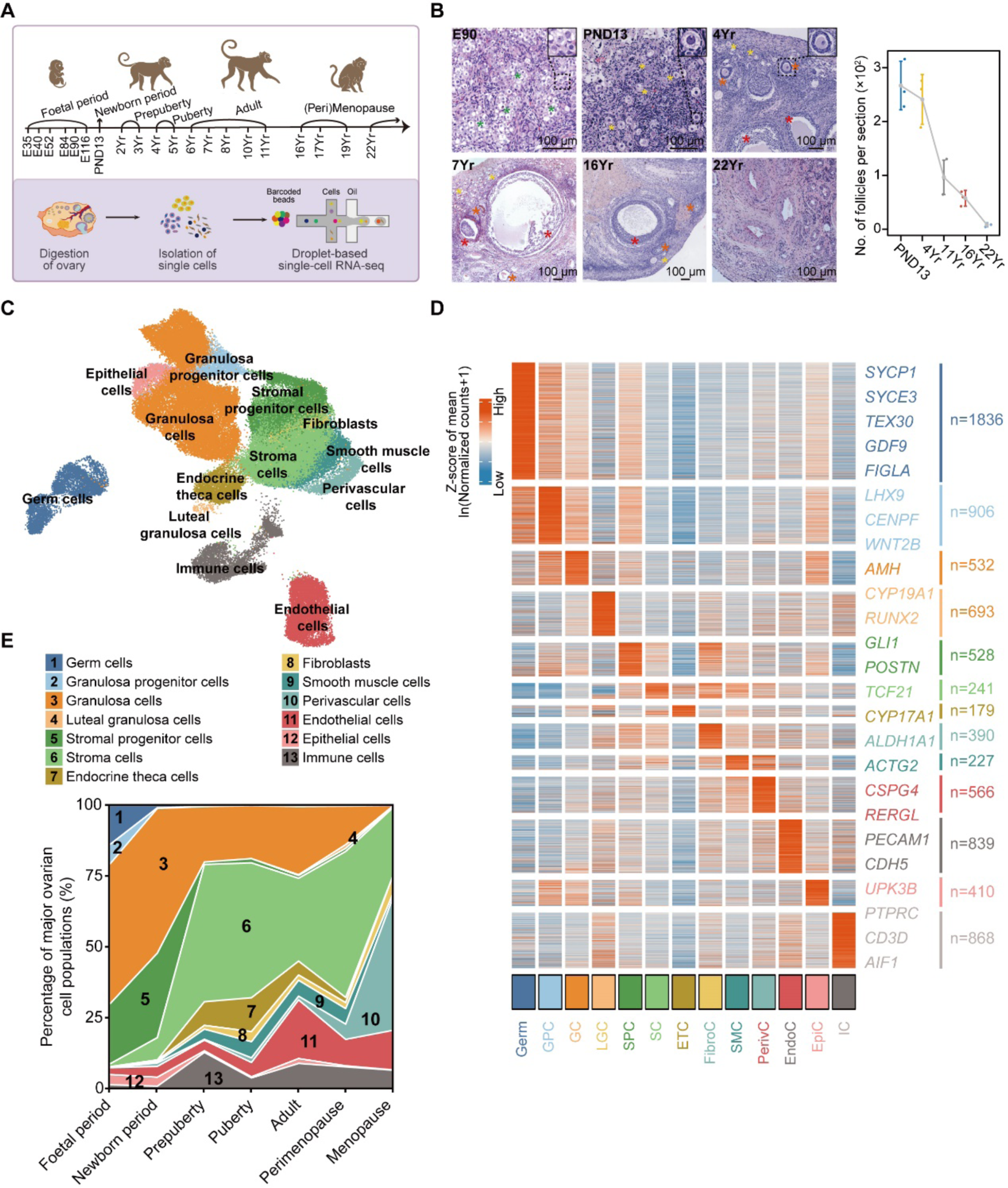
Cell atlas of cynomolgus monkey ovaries in the foetal, newborn, prepubertal, pubertal, adult, perimenopausal and menopausal stages. **A,** Single-cell RNA-seq workflow of ovarian cells of cynomolgus monkeys (*Macaca fascicularis*). The ages of the individual monkeys included in this study are indicated. E, PND, and Yr denote embryonic days, postnatal days and years, respectively. **B,** Left panel: H&E staining of ovarian tissues of cynomolgus monkeys at the indicated ages. Green stars indicate oocytes ahead of follicle assembly; yellow stars indicate nongrowing follicles; orange stars indicate growing follicles; red stars indicate antral follicles. The morphology of representative oocytes (framed by dashed lines) in ovaries of the corresponding ages are indicated, with magnified images in the top right corner. Scale bars are as indicated. Right panel: Measurement of follicle density in ovaries of different ages. Take sections of each ovarian sample with an interval of more than 1 mm and count the number of follicles (n=3). **C,** UMAP embedding visualization for 86,699 single cells from cynomolgus monkey ovarian tissues across all stages. **D,** Heatmap showing representative DEGs identified in different cell populations. Gene expression levels are averaged and scaled. Cell populations are colour coded as in (**C**). Germ, GPC, GC, LGC, SPC, SC, ETC, FibroC, SMC, PerivC, EndoC, EpiC and IC denote germ cells, granulosa progenitor cells, granulosa cells, luteal granulosa cells, stromal progenitor cells, stromal cells, endocrine theca cells, fibroblasts, smooth muscle cells, perivascular cells, endothelial cells, epithelial cells and immune cells, respectively. **E,** Stacked area plot showing the fraction of major ovarian cell populations in cynomolgus monkeys during the developmental stages and aging.

A total of 86,699 single cells passed stringent quality control measures for the downstream analysis (Fig. 1, D and E). We identified thirteen major populations by enriched gene expression profiles and signature genes (*35–42*) (Fig. 1, C and D and fig. S1F), containing all expected cell populations in the ovary (Fig. 1D), as reported previously (*15, 26–28, 43*). The cell populations and constitutions exhibited notable changes during monkey development, especially when entering the newborn, prepubertal and menopausal stages (Fig. 1E and fig. S2). The proportion of oocytes decreased with age, while fibroblasts and smooth muscle cells remained consistent throughout all stages (Fig. 1E and fig. S2). More importantly, we identified granulosa progenitor, precursor granulosa (pre-granulosa) and stromal progenitor cells, and these cells could be abundantly detected only in the foetal and newborn stages (Fig. 1C to E and fig. S2).

### Identification of PF assembly oocytes and their transcriptomic features

The formation of PFs is a critical step not only for the maturation of oocytes but also for the ovarian reserve in females (*44, 45*). Although the development of primordial germ cells has been studied in humans (*46*), the transition from these foetal germ cells (FGCs) to PFs is still unknown due to the ethical issues of using late gestation human embryos for research. As we obtained samples covering the entire ovarian developmental process in cynomolgus monkeys, we first analysed the FGCs and oocytes that were detected in our datasets.

We examined clusters of oocytes and identified seven subtypes reported in either humans (*46*) or monkeys (*28*): primordial germ cells, retinoic acid (RA)-responsive FGCs, meiotic oogonia (1 and 2), PF assembly oocytes, primordial follicle oocytes and two groups of growing follicle oocytes (oocytes 1 and oocytes 2). Meanwhile, the velocity cell trajectory analysis also revealed that the distribution of oocytes of different subtypes followed the expected pattern of oocyte maturation (Fig. 2, A and B and fig. S3A). These subpopulations were further validated by immunofluorescence staining of key marker genes that were identified in scRNA-seq data (Fig. S3, B and C). Interestingly, a unique group of oocytes was detected in the foetal stages and disappeared in the newborn samples (Fig. 2, A to C). These oocytes highly expressed typical genes, such as FIGLA and RHOXF1 (Fig. 2B and fig. S3A). Unlike the majority of oocytes of foetal ovaries (E90), which were closely associated in clusters (fig. S3B), the RHOXF1-expressing oocytes showed nest breakdown and became enveloped by a single layer of GCs from the surrounding nest (Fig. 2C). Moreover, signals for RHOXF1 could hardly be detected in PFs from the newborn ovaries (Fig. 2C); thus, we termed the RHOXF1-positive oocytes PF assembly oocytes. We further counted the RHOXF1 oocytes in E90 sectioned ovaries and found that these oocytes accounted for 9.66% of all oocyte population (Fig. 2C and fig. S3D), indicating the beginning of the PF pool establishment at this stage.

**Fig. 2.**
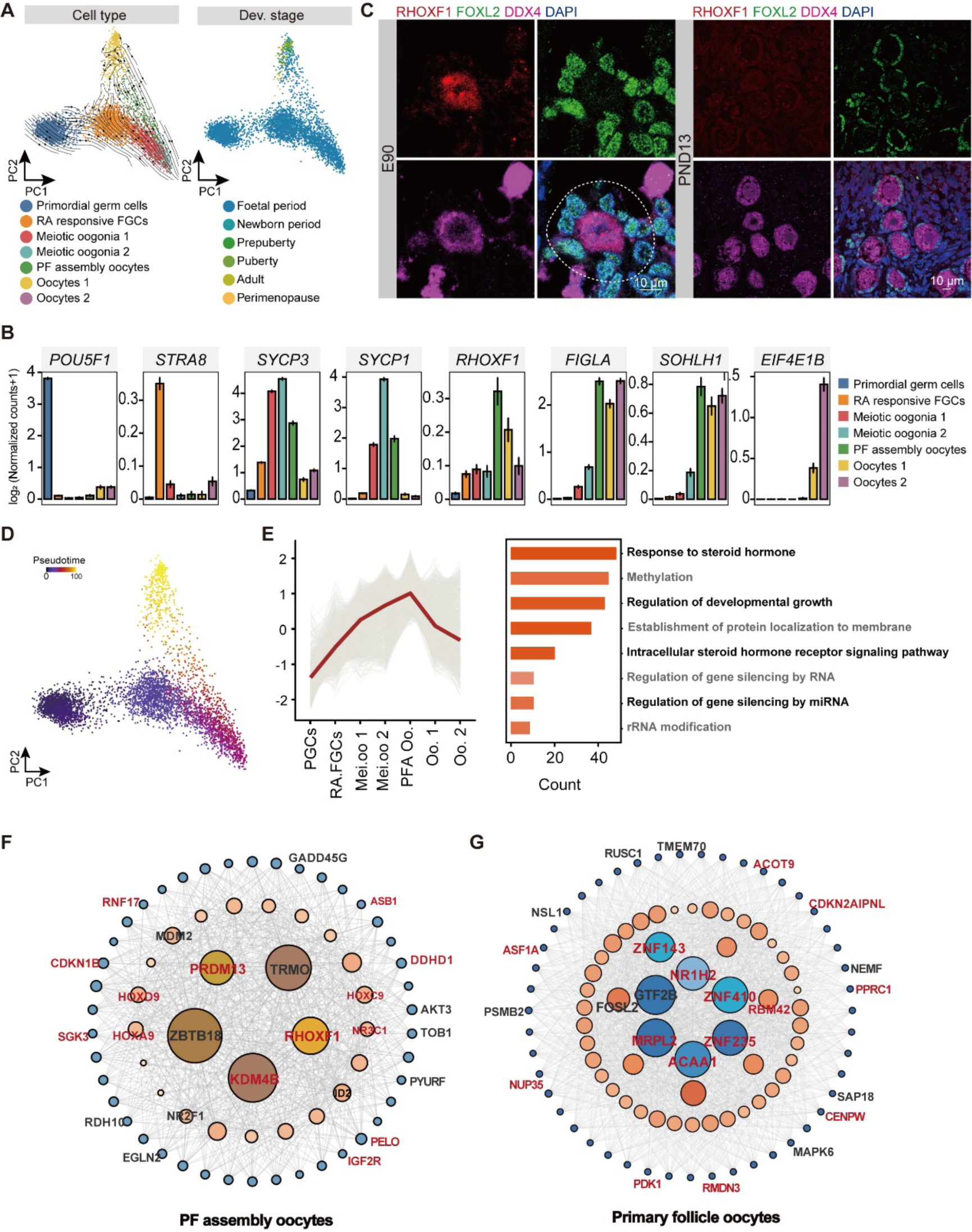
Transcription signatures of oocytes during PF assembly in cynomolgus monkeys. **A,** RNA velocity on PCA embedding visualization for seven subpopulations in germ cells identified in Fig. 1c. Cells are colour coded by the subpopulations identified (left) and developmental stage for cynomolgus monkeys where cells were from (right). RA denotes retinoic acid; FGCs denotes foetal germ cells, and PF denotes primordial follicles. **B,** Bar plots showing the expression levels of representative genes that are explicitly expressed in the indicated subpopulation. RA denotes retinoic acid; FGCs denotes foetal germ cells, and PF denotes primordial follicle. The error bars indicate mean values and standard error. **C,** Immunofluorescence staining for RHOXF1 (marker of PFA Oo), FOXL2 (marker of GCs, hereinafter inclusive) and DDX4 (marker of oocytes, hereinafter inclusive) in ovarian tissues from E90 and PND13 cynomolgus monkeys. The nuclei were counterstained with DAPI. The dashed line indicates the follicle during assembly. Scale bars are as indicated. **D,** Pseudotime trajectory of germ cells in Fig. 1C. Cells are colour coded by pseudotime inferred in the trajectory analysis. **E,** Representative gene cluster identified in PFA oocytes. Genes were clustered by their expression pattern along the developmental stage of germ cells by using the R package Mfuzz. Representative GO terms for these genes are shown on the right. PGCs denotes primordial germ cells; FGCs denotes foetal germ cells; RA denotes retinoic acid; Mei. oo denotes meiotic oogonia; PFA Oo denotes primordial follicle assembly oocytes; Oo denotes oocytes. **F,** Representative regulatory network identified in PF assembly oocytes. Key transcriptional regulons are represented as large coloured circles, and corresponding target genes are represented as small coloured circles. The node size indicates the number of target genes. Representative regulons and their target genes are highlighted. PF denotes the primordial follicle. Partial genes were retrieved from mouse ovarian scRNA data. Genes in black represent conserved genes expressed in both monkeys and mice, while genes in red represent monkey-specific expressed genes. **G,** Representative regulatory network identified in primary follicle oocytes. Key transcriptional regulons are represented as large coloured circles, and corresponding target genes are represented as small coloured circles. The node size indicates the number of target genes. Representative regulons and their target genes are highlighted. Partial genes were retrieved from mouse ovarian scRNA data. Genes in black represent conserved genes expressed in both monkeys and mice, while genes in red represent monkey-specific expressed genes.

We further examined the transcriptional signatures of PF assembly oocytes and reconstructed the pseudotime trajectory based on our scRNA-seq data (Fig. 2, D and E). Consistent with the cell identity, we clearly found that oocytes were distributed along the trajectory in a developmental time order (Fig. 2, D and E). To explore the temporal dynamics of the oocyte transcriptome, we clustered genes based on their expression patterns, identified seven time-dependent patterns and investigated their biological processes. Gene Ontology (GO) analysis showed that these genes were associated with certain biological functions corresponding to relevant stages (Fig. 2E and fig. S3, E and F). In particular, genes for PF assembly oocytes were significantly associated with “Response to steroid hormone”, “Methylation”, and “Regulation of developmental growth” (Fig. 2E). These temporally expressed genes included canonical (*FIGLA*, *JAG1*, *NOBOX*) and newly identified genes, such as *RMND5A*, *MAP1LC3A*, *BNIP3* and *YBX3* (fig. 3F). We also identified potentially important transcription factors (TFs) that participate in each stage. *RHOXF1* was identified as a pivotal TF in PFA oocytes by using the pySCENIC algorithm, and its major targets included *AKT3*, *TOB1*, *PYURF,* which are also conservatively expressed in mice. Furthermore, we observed other target genes such as *PRDM13* and *KDM4B* specifically expressed in cynomolgus monkey ovary (Fig. 2F). Interestingly, *RHOXF1* is conserved between humans and monkeys but the homologous gene (*Rhox10*) only expresses in mice testis, indicating its specific involvement in the regulation of the oocyte transcriptomic network during PF assembly in primates. In addition, we identified important TFs in primary follicle oocytes, of which *FOSL2* and *GTF2B* have been well studied previously (Fig. 2G), suggesting the involvement of these factors in PF activation and follicle growth in primates.

**Fig. 3.**
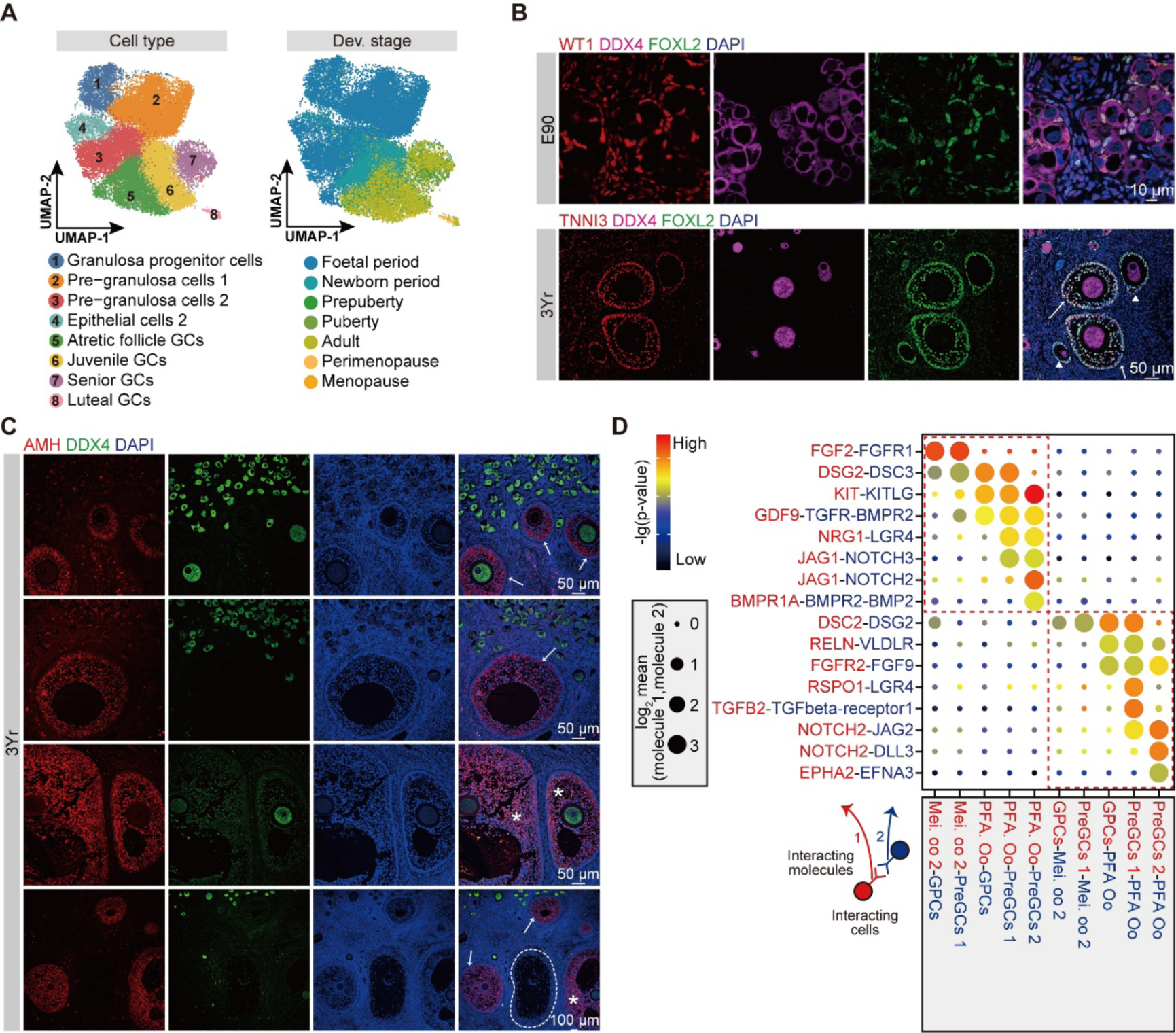
Constitution of GCs and their interaction with oocytes. **A,** UMAP embedding visualization for eight subpopulations in GCs identified in Fig. 1C. Cells are colour coded by the subpopulations identified (left) and developmental stage for cynomolgus monkeys where cells were from (right). GCs denotes granulosa cells. **B,** Immunofluorescence staining for WT1 (marker of granulosa precursor cells, hereinafter inclusive), DDX4 and FOXL2 in ovarian tissues from E90 cynomolgus monkeys and TNNI3 (characterized GCs from primary to antral follicles, hereinafter inclusive), DDX4 and FOXL2 in ovarian tissues from 3 Yr cynomolgus monkeys. The nuclei were counterstained with DAPI. The white arrowheads point to the primary follicles, and the arrows show the secondary follicles. Scale bars are as indicated. **C,** Immunofluorescence staining for AMH (marker of GCs from secondary and antral follicles, hereinafter inclusive) and DDX4 in ovarian tissues from 3 Yr cynomolgus monkeys. The nuclei were counterstained with DAPI. White arrows point to secondary follicles, asterisks indicate antral follicles, and dashed circles delineate atretic follicles. Scale bars are as indicated. **D,** Dot plot depicting representative ligand-receptor interactions between oocytes and granulosa cells in different development stages. Interaction strength was measured using log2-scaled means of the average expression level of the ligand in the indicated cell type and receptor in the other cell type. The p values (permutation test) are colour coded. GPCs, PreGCs, Mei. oo, PFA denote granulosa progenitor cells, pre-granulosa cells, meiotic oogonia and primordial follicle assembly, respectively.

### The precursors and differentiation of GCs in primates

GCs are pivotal for ovarian follicle assembly and growth through crosstalk with oocytes (*13, 47–50*). However, the differentiation of GCs and their interplay with oocytes are poorly characterized in primates. We found that the GC populations could be divided into seven subtypes according to their gene expression patterns (Fig. 3A, fig. 4 A to C). Besides, it is reported GCs are partially originated from epithelial cells in rodents (*47*). Here we found the epithelial cells could be clustered into two subtypes: epithelial cells 1 appeared from foetal period and expressed traditional epithelial makers (fig. S4, C and E); epithelial cells 2 were abundant in the early ovaries (E38-PND13) and expressed granulosa progenitor cell markers (WT1, RSPO1, LHX9) (Fig. 3A, fig. S4D). Specifically, the granulosa progenitor cells gave rise to the precursor GCs (I: corresponding to the first waves of primordial follicles; II: corresponding to the second wave of primordial follicles and primary follicles) and could barely be detected after birth (fig. S2, fig. S4, A and B); notably, the epithelial cells 2 persistent from foetal to newborn period, suggested they provide precursor GCs continuously for the establishment of primordial follicle pool (Fig. 3A, fig. S2, fig. S4, A and B); the juvenile GCs (corresponding to secondary follicles) appeared sporadically at foetal period, and rapidly increased from newborn period; correspondingly, the number of atretic follicle GCs increased as well; the senior GCs (corresponding to antral follicles) could be steadily detected from prepuberty to perimenopause; the luteal GCs arose from puberty, indicated ovulation and menstruation (Fig. 3A, fig. S2, fig. S4, A and B). However, in menopause, GCs were scarcely detected in monkey ovaries (fig. S2).

**Fig. 4.**
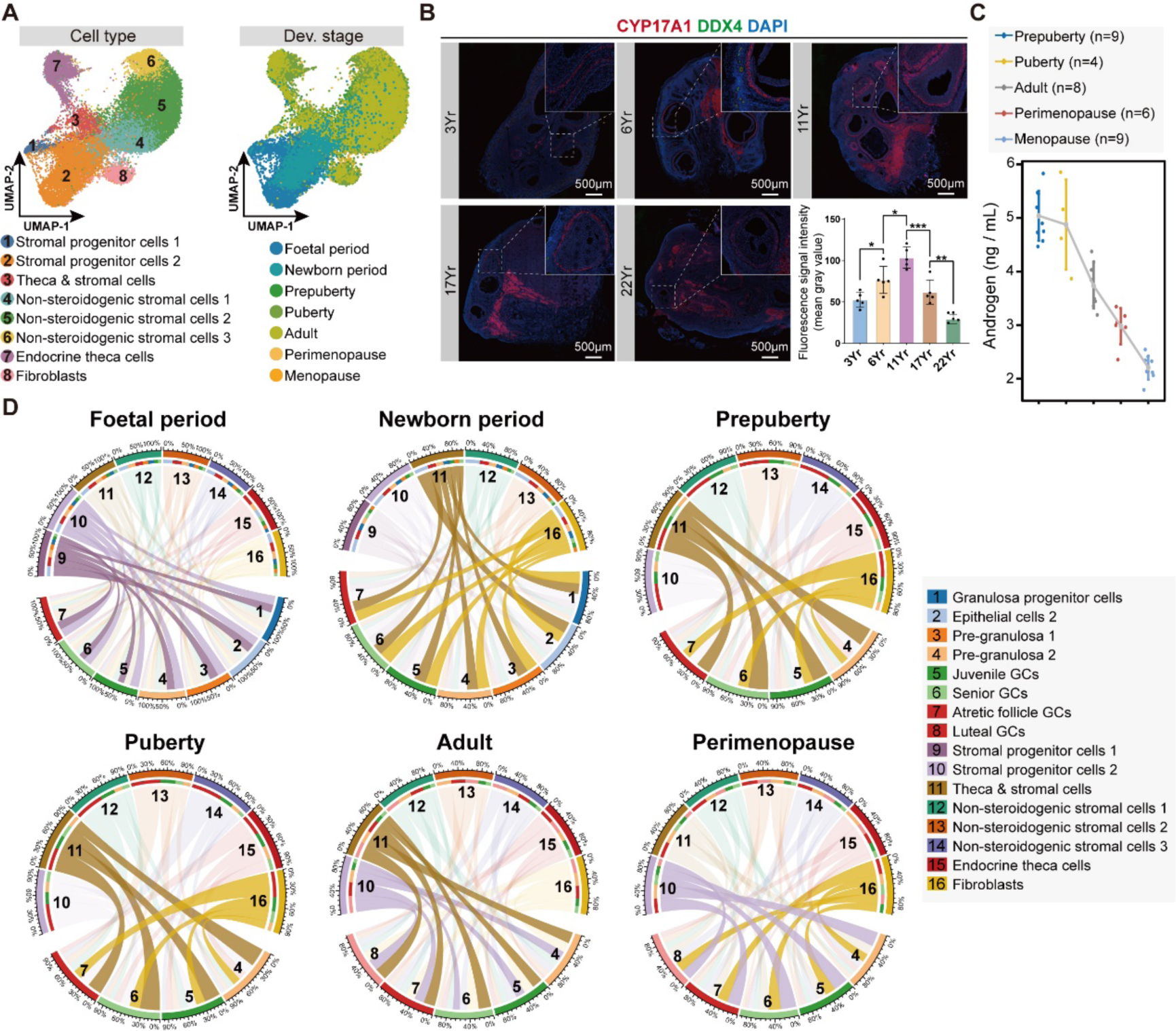
Analysis of stromal cells during cynomolgus monkey development and their interaction with GCs. **A,** UMAP embedding visualization for eight subpopulations in stromal progenitor cells, stromal cells, endocrine theca cells, and fibroblasts identified in Fig. 1C. Cells are colour coded by the subpopulations identified (left) and developmental stage for cynomolgus monkeys where cells were from (right). **B,** Immunofluorescence staining for CYP17A1 (marker of theca cells), DDX4 in ovarian tissues from cynomolgus monkeys and statistical analysis of the expression intensity of CYP17A1 indicated by red fluorescence. The nuclei were counterstained with DAPI. The dashed frames indicate theca cell layers, and magnified images are on the upper right corner. The age of individual monkeys and the scale bars are as indicated. n=3 sections, *p < 0.05, **p < 0.01 and ***p < 0.001. **C,** The variation in androgen concentrations in the peripheral blood of cynomolgus monkeys at different age stages. In the error bar plots, mean±SE was used for visualizing errors. **D,** Chord diagrams showing interactions among subpopulations in GCs, stromal progenitor cells, stromal cells, endocrine theca cells and fibroblasts in Fig. 1C in foetal period, newborn period, prepuberty, puberty, adult, and perimenopause. GCs denotes granulosa cells.

Furthermore, we defined differentially expressed genes (DEGs) among subpopulations in GCs and confirmed representative DEGs in each group by immunofluorescence staining. WT1 (*51*), the marker for granulosa progenitor cells, could be abundantly detected during the foetal period (Fig. 3B, fig. S4, C, D and F). For AMH highly expressed in juvenile and senior GCs, immunofluorescence staining further confirmed its expression in secondary and antral follicles but not in primary and atretic follicles (Fig. 3C and fig. 4G). Consistently, the GC marker TNNI3 was highly expressed from secondary to antral follicles and sparsely expressed in some primary follicles (Fig. 3B and fig. S4H). INHBB was identified as the marker for senior GCs (fig. S4, C and D) and was detected exclusively in GCs of the antral follicles (fig. S4I).

To explore communication between oocytes and surrounding GCs, we measured cell-cell interactions at corresponding stages by using CellPhoneDB. The results demonstrated that FGF2-FGFR1 was the exceedingly active ligand-receptor interaction at meiosis, while KIT-KITLG and JAG-NOTCH were the most significant interactions during PF assembly (Fig. 3D). Previous reports showed that macrophages are involved in intraovarian events (*52–56*), especially the elimination of apoptotic GCs from atretic follicles. To elucidate the roles of immune cells in ovarian development, we grouped and characterized these cells into three subtypes, macrophages, T lymphocytes and B lymphocytes, and the macrophages were obviously the largest group (fig. S5, A to C). The characteristic genes of macrophages and T lymphocytes were validated by immunofluorescence staining (fig. S5D). Intriguingly, when we examined the interplay between GCs and macrophages from foetal period to perimenopause (fig. S5E), we found that all GC subgroups represented interactions with macrophages, indicating that the molecules that connect macrophages were already prepared at the mRNA level in GC populations before apoptosis.

### In-depth analysis of somatic cells revealed their functions during ovarian development of cynomolgus monkeys

Stromal cells are the most abundant cell populations in the ovary. However, the classification and the developmental trajectory of this major class are exceedingly poorly studied in primates (*43*). To fill this gap, we explored and identified eight subpopulations of stromal cells in depth (Fig. 4A, fig. S6 A to C). We found two stromal progenitor clusters, the first cluster was mainly found in early foetal period and strongly expressed MKI67, indicated they were proliferative (Fig. 4a and fig. S6 A to C); the stromal progenitor 2 were validated by POSTN staining, they were substantially existed in foetal and newborn periods (fig. S6D). Theca & stromal cells may serve as precursors of endocrine theca cells and nonsteroidogenic stromal cells (Fig. 4A) and consecutively exist from the foetal period to perimenopause (fig. S2 and fig. S6A). In particular, the stromal progenitor cells 2 might also derive a population of fibroblasts that has not been specified before. The fibroblasts highly expressed genes for extracellular matrix (ECM) organization (fig. S6, B and C) and could be readily detected from foetal period to menopause, suggesting essential roles in ovarian organization (Fig. 4A and fig. S2). Endocrine theca cells are pivotal for follicle development due to androgen secretion (*57, 58*). CYP17A1 and STC1 are markers for this population were also validated by immunostaining (Fig. 4B and fig. S6E).

Endocrine theca cells were firstly appeared in foetal period and abundantly from prepuberty to adult, and the proportion decreased markedly from the adult stage to menopause (fig. S2 and fig. S6A). Correspondingly, the androgen levels in peripheral blood throughout the cynomolgus lifespan were constantly reduced from puberty to menopause, indicating the dysregulation of this hormone (Fig. 4C). Furthermore, the nonsteroidogenic stromal cells could group into 3 subpopulations, and there were drastic changes in cell proportions from foetal period to menopause (fig. S2 and fig. S6 F to G). In addition, we investigated the interplay between stromal cells and GCs. Stromal progenitor cells have the most significant correlations with GCs in fetus, then after birth, theca & stromal cells and fibroblasts take place to communicate most with GCs until adulthood period. Theca & stromal cells have the most frequent communication with GCs from newborn to adulthood stages (Fig. 4D). At perimenopause, fibroblasts showed the greatest communication with GCs (Fig. 4D). Moreover, we measured the crosstalk between stromal lineages and macrophages. We found the non-steroidogenic stromal cells 1/2 had strong connections with macrophages at foetal and newborn stages, which weakens after birth, particularly in case of stromal cells 2. The theca & stromal cells and fibroblasts had strong interaction with macrophages across lifespan (fig. S6H).

To explore the cellular diversity and their trend during ovarian development, we also studied other somatic cells in the ovary. The proportion of endothelial cells existed steadily throughout the ovarian life course (fig. S2 and fig. S7A). The results showed that endothelial cells were classified into 3 groups, with different signature genes (fig. S7, A and B). Smooth muscle cells and perivascular cells have been reported to originate from a common mesenchymal progenitor (*59*). Here, we demonstrated that perivascular cells could be distinguished from smooth muscle cells (fig. S7C). ACTA2 could be detected in both cell types, while CSPG4 was almost specifically expressed in perivascular cells (fig. S7D). In addition, endothelial, smooth muscle and perivascular cells were previously reported to engage in angiogenesis. Here, we found that the proportions of perivascular cells remarkably increased starting from perimenopause (Fig. 1E and fig. S2). Consistent with this finding, the level of vascularization increased gradually to a plateau at menopause, consistent with the findings of another study (*60*) (fig. S7, E and F).

### Cell type-specific transcriptomic clocks quantified primates biological ovarian aging

In contrast to sparsely obtained ovarian tissues mainly from adult human patients, our monkey ovarian atlas offers the opportunity to establish a precise molecular clock for the evaluation of ovarian aging and female fertility. We first selected stromal, vascular, immune, and granulosa cells as speculated they might effectively predict the ovarian biological age due to their significant changes in transcriptomes and cell compositions (Fig. 1E and fig. S2). To achieve this, we utilized the gene expression profiles of ovarian cell and introduced a LASSO regression model (*61*) to train the single-cell transcriptomic data for developing aging clocks. The clocks can predict the ovarian biological age with correlations (R) greater than 0.80, and the stromal cells’ clock got the most accurate prediction (R=0.91) (fig. S8A and fig. S9). To pinpoint which cell type is most predictive of the ovarian aging, we trained the transcriptomic data of all the 17 ovarian cell types by using the regression model and developed 17 cell type-specific aging clocks (Fig. 5A and fig. S8B). The clocks could predict biological ovarian aging with high accuracy, especially constructed from juvenile/senior GCs, epithelial cells 1, and non-steroidogenic stromal cells (Fig. 5A). It is notable that the observed aging clock genes are predominantly cell-specific, and display minimal inter-gene overlap (Fig. 5B). In order to obtain a more thorough comprehension of the functional modifications associated with ovarian aging in a cell-specific manner, we have provided an account of changes in the enrichment of cell type-specific gene ontology (GO) during the aging process. The examination revealed a range of GO terms that display a consistent trend of enrichment levels with age in specific cells (Fig. 5C and fig. S10). For instance, age-associated alterations in the functionality of juvenile GCs manifest as “response to corticosteroid” and “reproductive structure development”. Non-steroidogenic stromal cells 1 exhibit significant functional differences in aspects such as “cell growth” and “response to steroid hormone” (Fig. 5C). To investigate the mechanism of how individual cells contribute to ovarian aging, we performed Kyoto Encyclopedia of Genes and Genomes (KEGG) pathway enrichment analysis on a cell-specific gene set that constructs the aging clock. The AGE-RAGE signaling pathway identified in juvenile GCs and the relaxin signaling pathway found in non-steroidogenic stromal cells 1 are considered to be the most prominent pathways. The violin plots depicting the expression of relevant genes in the pathways also illustrate unique trends across different stages (Fig. 5D).

**Fig. 5.**
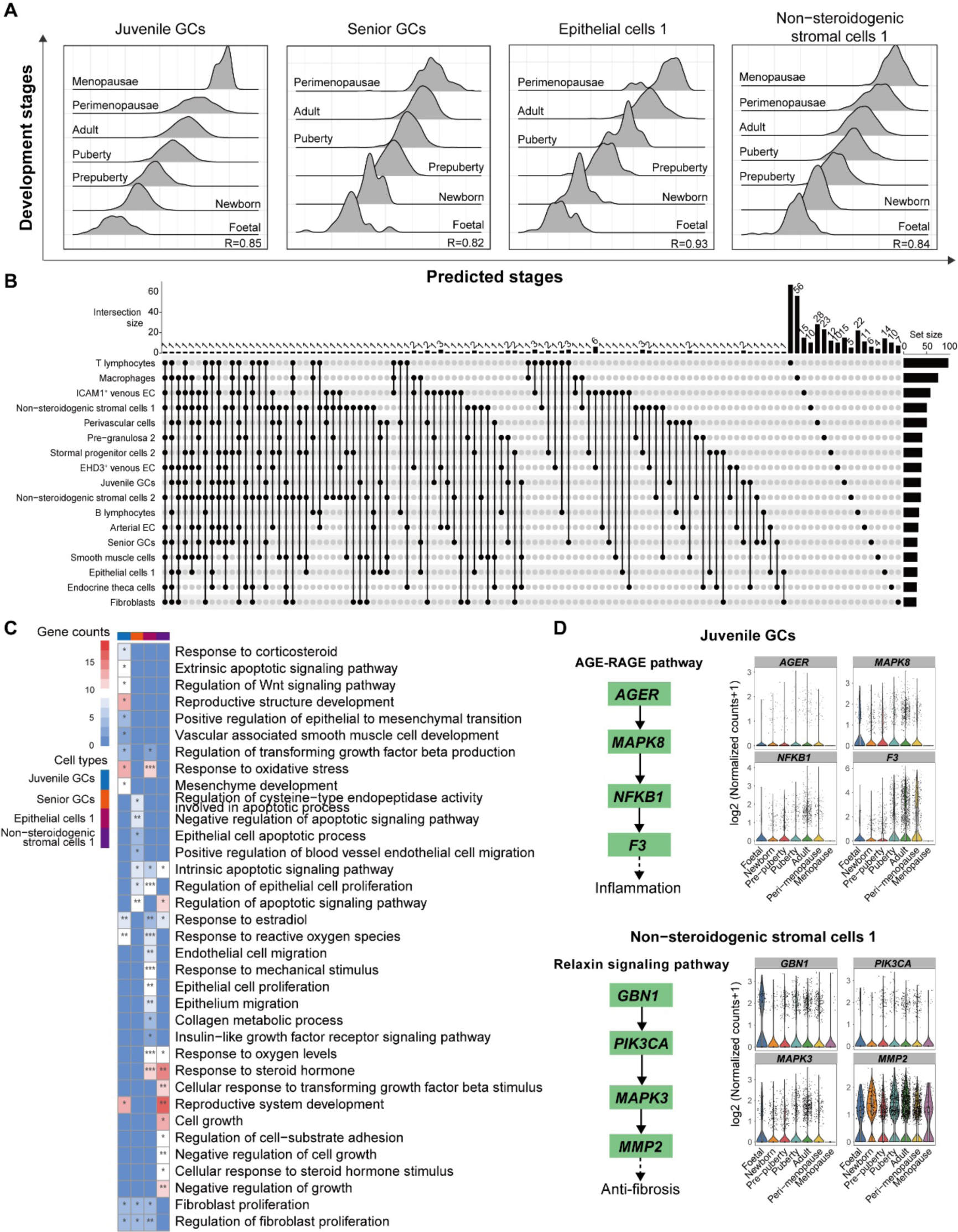
Construction of ovarian cell-type-specific aging clocks. **A,** Ridge plots showing associations between developmental stages and predicted stages based on cell-type-specific aging clocks. GCs denote granulosa cells. The Pearson’s correlation coefficient (R) were as indicated. **B,** UpSet plot showing the intersection of genes used by cell-type-specific aging clocks. ECs denotes endothelial cells and GCs denotes granulosa cells. **C,** Heatmap showing representative GO terms for genes used in cell type-specific clocks. GCs denotes granulosa cells. *p < 0.05, **p < 0.01 and ***p < 0.001. **D**, Violin plots showing expression levels for genes involved in AGE-RAGE pathway and Relaxin signaling pathway. GCs denotes granulosa cells.

To investigate the correlation between the aging rate of specific cell type and the whole ovary, we predicted the biological age of the entire ovary represented all the cells, and compared it to individual cell types (the slope of the line [k] represents the responsiveness to aging; “ Δk” represents the different degrees of response to aging, where a negative value indicates a deceleration and a positive value indicates an acceleration) (Fig. 6A). The results showed that the aging rate of single cell type is delayed compared to the rate of overall ovary, with the aging rate of fibroblasts, smooth muscle and epithelial cells 1 being the closest to the overall rate of ovarian aging (Fig. 6A and fig. S8C).

**Fig. 6.**
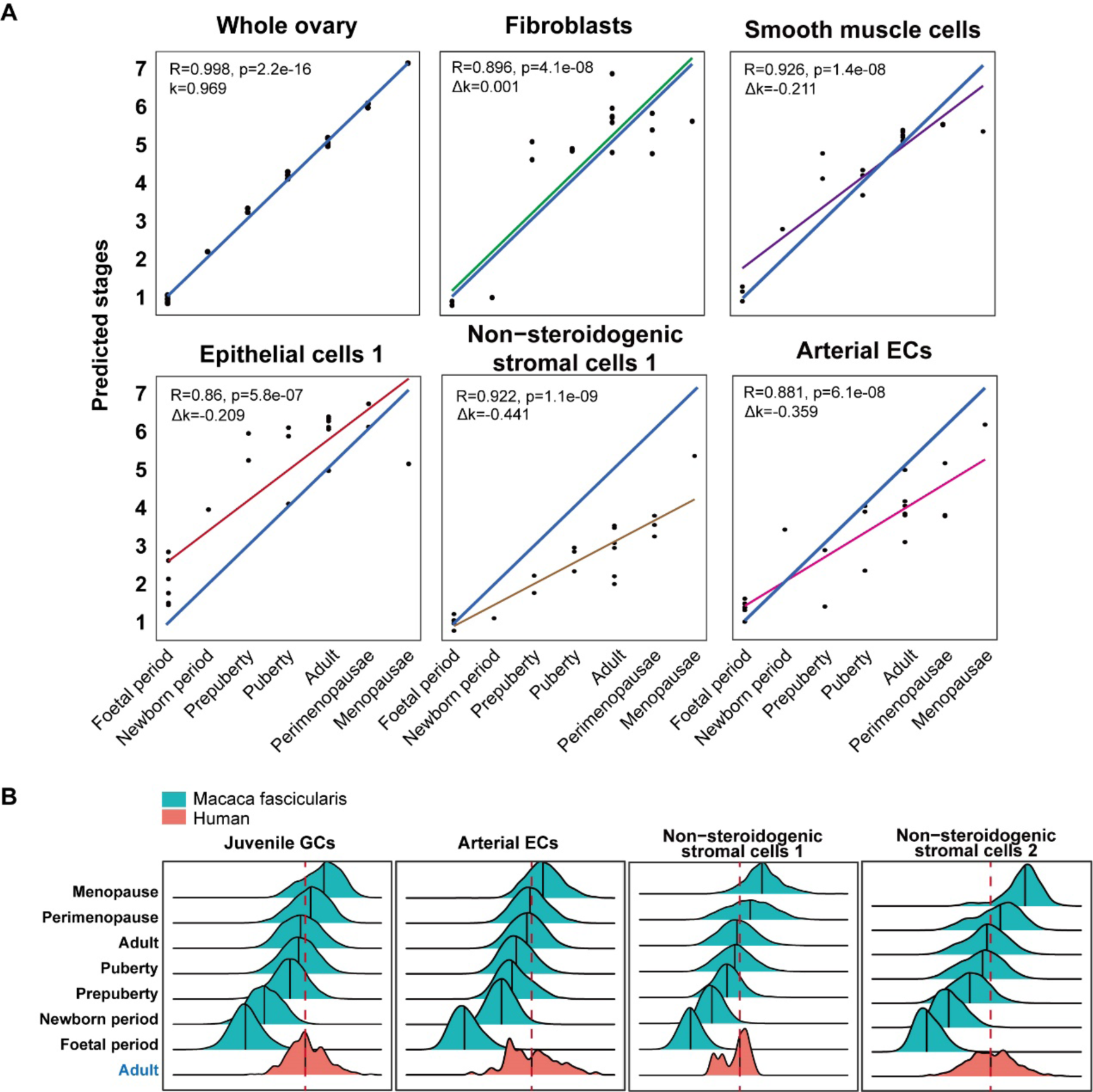
The accuracy of cell-type-specific aging clocks in characterizing ovarian aging and its validation in human. **A,** Scatter plot showing correlations for average expression levels of whole ovary and a specific cell type. The R value indicated correlation coefficient, k value represents the responsiveness to aging and delta k (Δk) indicated the difference of response to aging between the individual cell types and the whole ovary. ECs denotes endothelial cells. The p value was as indicated. **B,** Ridge plots showing associations between developmental stages and predicted stages based on cell-type-specific aging clocks. GCs denotes granulosa cells. ECs denotes endothelial cells. The dashed lines indicate the median value.

### The ovarian clock was conserved between humans and monkeys

NHPs have long been used as surrogates for humans due to their close evolutionary relationship with human beings (*28, 62*). However, the homogeneity between the two species in ovarian cells is unknown. To determine the conservation of diversity in cell populations, we performed integrated analysis with datasets from adult human ovaries (fig. S11 and fig. S12). The comparison revealed well-conserved cell populations that enable matching of cell types of cynomolgus monkeys to human adults (fig. S11, A and B). Significantly, we identified two more crucial cell populations in monkeys, granulosa precursor and stromal progenitor cells, benefitting from the comprehensive sampling of cynomolgus monkeys (fig. S11, A and B). To systematically compare monkey and human ovarian cells, we further matched all ovarian cell subpopulations identified in adult monkeys. All subpopulations exhibited general consistency. The proportions of the majority of subpopulations in the two species were comparable (fig. S11C).

Notably, the detailed classification and validation of ovarian cells of cynomolgus monkeys in this study offered the opportunity to refine the subgroups of ovarian cells in human adults, which uncovered those previously undefined cell types in humans (*26*), including perivascular cells and subclusters of stromal and endothelial cells. To further elucidate the convergent and divergent features of the two species, we calculated the correlation coefficients of all ovarian somatic cells and found a comprehensive similarity score (fig. S12A). Notably, most of the ovarian cell types were conserved between the two species except the luteal GCs (fig. S12A). Furthermore, we defined 1,553 DEGs from all ovarian somatic subpopulations of cynomolgus monkeys and found that these genes could distinguish subpopulations in humans (fig. S12B) and vice versa (fig. S12C). Thus, the integrated analysis revealed a well-conserved cellular map and highlighted the merits of our data for gaining insights into human ovarian development, aging and dysfunction.

The above investigation prompted us to learn whether there are inter-primate differences when it comes to transcriptomic age of ovaries. To externally validate these cell type-specific aging clocks, we retrained the monkey ovarian clocks and applied them to independent human datasets (*26*). The results showed that monkey ovarian clocks could predict chronological age accurately of juvenile GCs and non-steroidogenic stromal cells 1/2 (Fig. 6B), suggesting broad applications of monkey ovarian aging clocks to quantify human samples and evaluate female fertility.

## Discussion

The underlying pathophysiology of age-related changes within the ovary remains poorly understood. Discovering the molecular mechanisms governing ovarian aging and identifying biomarkers which could help identify women at increased risk of accelerated reproductive senescence may permit timely preventative action and critical for contemporary reproductive research. During the past few years, advanced single-cell technologies have allowed elucidation of the cellular constitutions and molecular features of human early embryos, which showed significant divergence between humans and mice in gonad development (*63*). However, the postnatal development of human ovaries is largely unknown due to ethical issues in acquiring healthy samples. The cross-species analysis of ovarian cells between monkeys and human adults showed that the cellular composition and gene expression are generally similar between these two species (fig. S11 and fig. S12), suggesting that cynomolgus monkeys are a good surrogate to study human ovary development and dysfunction. More importantly, our study provided a long-term course of ovarian cell dynamics in primates through the analysis of tens of thousands of single-cell transcriptomes, which allowed us to decipher ovarian development and cell-cell interactions along temporal trajectories. The 22 samples used in this study span 22 years, from the foetal period to menopausal stage, covering 7 pivotal stages during ovarian development. We derived a detailed transcriptional profile of 86,699 single cells and elucidated the developmental landscapes of the cynomolgus monkey. The datasets allowed observation of PF assembly and the interaction between somatic cells and somatic cells-to-oocytes. We observed PF assembly during the foetal period and identified ROHXF1 as a marker for these oocytes.

Furthermore, we displayed the dynamic diversification of the proportions of ovarian cells and found two populations of cells (granulosa progenitor cells and stromal progenitor cells) that have not been previously characterized in primates (Fig. 1 and fig. S2). According to our findings, granulosa precursor cells and stromal progenitor cells existed abundantly during the foetal period. Then, after birth, the granulosa precursor cells differentiated to the granulosa lineage; the stromal progenitor cells gave rise to the theca and the stroma lineages. Immune cells and stromal lineages are fundamental for folliculogenesis and ovarian development. In particular, it has been proved a group of epithelial pregranulosa (EPG) cells in the surface epithelium of murine ovary, which could further differentiate into granulosa cell of the primordial follicle, therefore participate in the establishment of primordial follicle pool (*47*). Here we identified two distinct epithelial cell types in foetal and newborn monkey ovary. Epithelial cells 2 express not only canonical epithelial markers (UPK3B, KRT19) but also genes associated with granulosa progenitor cells (WT1, RSPO1, LHX9), indicates this particular epithelial cells 2 predisposed towards the granulosa cell lineage in primates. Furthermore, folliculogenesis and ovarian development require the collaboration and interaction of multiple cell types, which are involved in perimenopause and ovarian senescence (Fig. 7). The findings also indicate that distinct transcriptional features may underlie cell-specific aging processes.

**Fig. 7.**
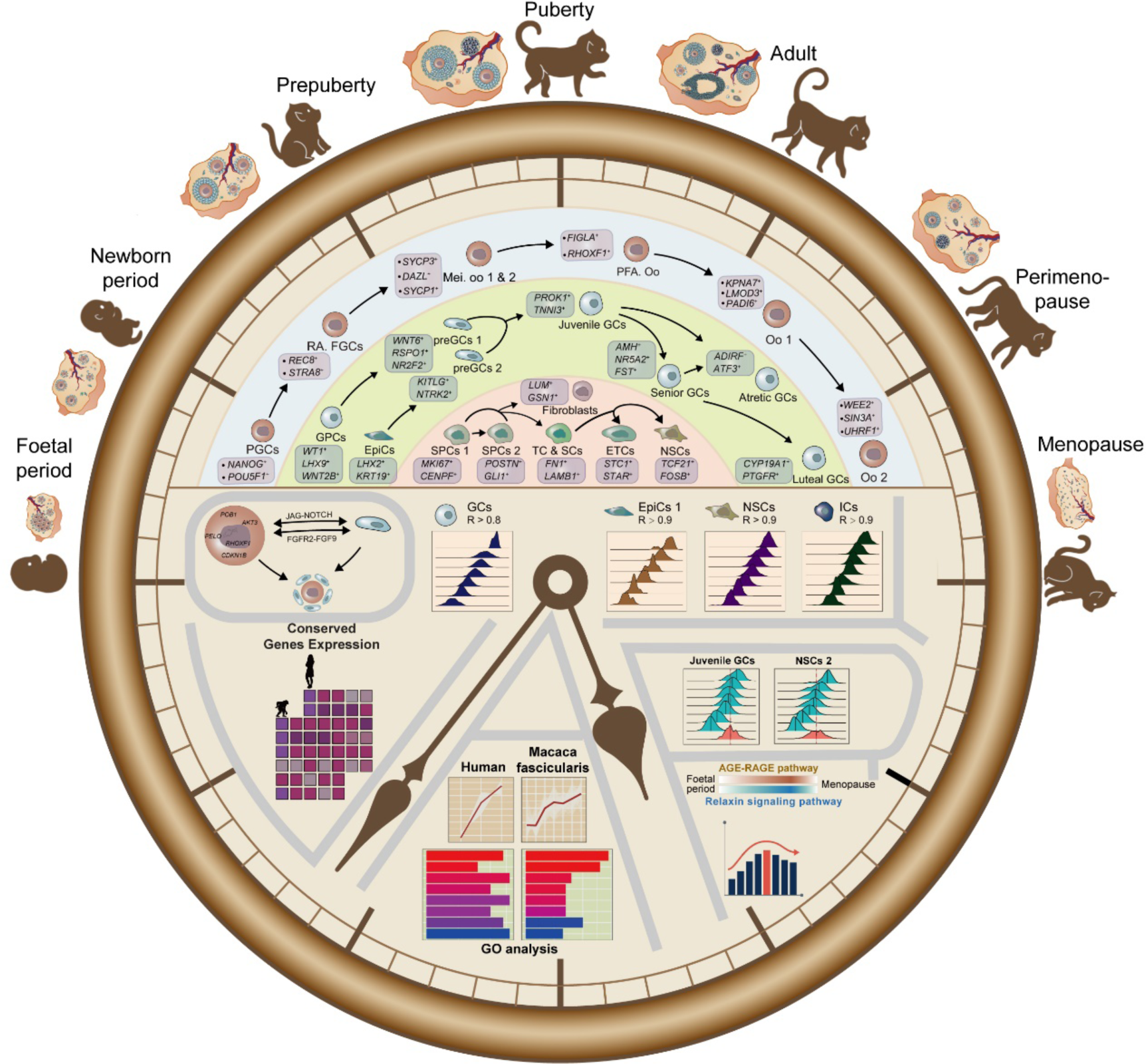
A conserved ovarian aging clock in primates based on longitudinal panorama of monkey ovary. The outer part of the schematic scheme indicates the change in morphology and the variety and order of internal cells of the ovary as it develops with age. The inner part of the schematic scheme indicates the developmental lineages of oocytes, GCs, theca cells and stromal cells and intercellular regulation of follicular genesis and development. Cell-type-specific fertility clocks established in cynomolgus monkeys could precisely predict the developmental stages for juvenile granulosa cells and non-steroidogenic stromal cells 2 in the human. The human traits were mapped to ovarian cells in cynomolgus monkeys. PGCs denotes primordial germ cells; RA denotes retinoic acid; FGCs denotes foetal germ cells; Mei. oo denotes meiotic oogonia; PFA denotes primordial follicle assembly; Oo denotes oocytes; GPCs denotes granulosa precursor cells; preGCs denotes pre-granulosa cells; GCs denotes granulosa cells; SPCs denotes stromal progenitor cells; TC denotes theca cells; SCs denotes stromal cells; ETCs denotes endothelial theca cells; NSCs denotes non-steroidogenic stromal cells; EpiCs denotes epithelial cells and ICs denotes immune cells.

Due to the poor accessibility of human organs, the samples utilized in previous studies are usually partial, diseased tissue and sparse in time points (*26, 27*). NHPs share similar genetic and physiological properties and represent the closest model for human research (*64*). With the advance of our monkey ovarian atlas, we identified and characterized the landscape of all cell types. This longitudinal study on independent cohort makes it possible to generate an accurate and biologically meaningful cell-type-specifically transcriptomic ovarian clock, which to our knowledge is the first report for building molecular clocks with ovarian single-cell transcriptomes in primates. The resulting predictor performs remarkably well across a wide spectrum of ovarian cell types, especially in granulosa, stromal, epithelial, endothelial, and immune cells. As it is easily to acquire these somatic cells as appendants of oocytes retrieval, this ovarian aging clock could conveniently predict the reproductive senescence and age-related health, especially for providing predictive value to identify patients at risk of subfertility. Patients at risk of earlier menopause could also benefit from lifestyle and medical interventions. Therefore, these ovarian clocks have the potential to be highly clinically used as prognostic and diagnostic markers of disease. Furthermore, we demonstrated the high-reliability of this clock by applying published human datasets in this algorithm.

Overall, our comprehensive analysis of the ovarian single-cell transcriptome throughout the monkey life cycle provides a rich foundation for understanding the function of ovarian cells and elucidating the regulatory insights underlying ovarian development, aging and dysfunction. The ovarian molecular atlas enabled us to depict cell-type specific ovarian clocks to predict biological ovarian aging. The macaque ovarian atlas would be an invaluable resource for human studies because of cellular homogeneity and limitations and the lack of availability of healthy human samples.

## Materials and methods

### Ethical statement

This study was conducted following the Principles for the Ethical Treatment of Non-Human Primates and was approved in advance by the Institutional Animal Care and Use Committee of the Institute of Zoology (Chinese Academy of Sciences, Appl. No: IOZ-IACUC-2021-174). All applicable institutional and/or national guidelines for the care and use of animals were followed.

### Collection of ovaries from cynomolgus monkeys

All 17 cynomolgus monkeys (*M. fascicularis*) were of Southeast Asian origin. The animals were kept at approximately 25 ℃ on a 12-hour light-12-hour dark schedule and raised at Beijing Institute of Xieerxin Biology Resource with the accreditation of the Laboratory Animal Care Facility in Beijing. None of the animals had a clinical or experimental history.

Cynomolgus monkeys were anaesthetized with ketamine (10–12 mg/kg) before surgery. The ovaries were obtained through surgical removal from the cynomolgus monkey. Nonovarian tissue was removed prior to processing. Then, the ovaries were washed thoroughly to remove the blood with sterilized precooled phosphate buffered saline (PBS). Subsequently, the ovaries were weighed and photographed. The ovaries were immediately transported to the laboratory on ice in medium containing α-MEM (Gibco, 12561-056), 1% HSA (Merck Millipore, 823022), 1% penicillin/streptomycin (Gibco, 15140-122). Finally, the ovaries were digested for single-cell sequencing or fixed with formalin for subsequent assays.

### Preparation of single-cell suspensions for scRNA-seq

Single-cell isolation was performed on the basis of a protocol described previously with modification. In brief, the “clean” ovaries were cut first into 1 mm thick pieces and then minced using a tissue sectioner (Mcllwain Tissue Chopper, The Mickle Laboratory, Guildford, UK). The minces were transferred to a 50 ml conical tube and digested enzymatically in a medium containing α-MEM (Gibco, 12561-056) + 0.04 mg/ml Liberase DH (Roche, 5401054001) + 0.4 mg/ml DNase I (Sigma, DN25) + 1% penicillin/streptomycin (Gibco, 15140-122) for 60 minutes in a shaker (160 rpm) at 37 ℃ interspersed with 20–30 pipetting every 15 minutes. The digestion was terminated by α-MEM media with 10% HSA (Merck Millipore, 823022). After digestion, the cell suspension was filtered by a 40-µm strainer and centrifuged at 300 × g for 5 minutes, followed by gentle removal of the supernatant. For removal of red blood cells, the cell pellet was treated with 3 ml of ACK (ammonium chloride-potassium) lysis buffer for 10 minutes at room temperature (RT), and then, the treatment was terminated with 27 ml of PBS. The remaining cells were washed twice with PBS and centrifuged at 300 × g for 5 minutes. For the single-cell suspension, the cell pellet was resuspended in PBS and adjusted to a concentration of 700-1,200 cells/μl to ensure subsequent cell capture efficiency. Cell concentration and viability were measured with a Countess II automated cell counter (Thermo Fisher, AMQAX1000). The cells were then used for single-cell sequencing.

### Single-cell RNA-seq library construction and sequencing

Cells were loaded onto a 10x Genomics Chromium chip targeting an 8,000-10,000 cell range as recommended by the factory. Reverse transcription and library preparation were performed using the 10x Genomics Single Cell v2 kit following the 10x Genomics protocol. The libraries were then run on an Illumina NovaSeq 6000 in two lanes to generate 150 PE reads.

### Preprocessing single-cell RNA sequencing data

Raw sequencing reads were processed using Cell Ranger (v3.1, 10x Genomics) with default parameters. The reference genome for read alignment and quantification was generated following instructions from 10x Genomics. The genome (Macaca fascicularis_5.0) and annotations were obtained from Ensemble (version 102). Briefly, raw reads were aligned with the reference genome and quality checked. Uniquely mapped reads were used for UMI counting. Gene expression levels were further quantified for each barcode observed. Cells with a much lower RNA content were discarded. The filtered feature barcode matrices generated by Cell Ranger (v3.1, 10x Genomics) were adopted for downstream analysis.

### Identification of cell populations in single-cell RNA sequencing data

Gene expression data were processed by using the R package Seurat (v4.0.4) (*65*). In this study, raw counts were scaled to 10,000, and expression levels were defined as log2 (normalized counts + 1). As a quality control, cells were filtered out if they failed to meet these criteria: 1) the number of detected UMIs was not less than 2,000; 2) the number of detected genes was not less than 800; and 3) the log-transformed ratio of detected genes and detected UMIs was higher than 0.8. Two rounds of clustering were performed. Consensus variable features among samples were first identified as previously reported. For individual samples, 3,000 features were selected. These selected features were merged, and a consensus list of 3,000 features was generated based on hits across samples. For removal of potential sample/batch effects, Scanorama (v1.7.1) was used for dimension reduction with these consensus features (*66*). After initial clustering, potential doublets in individual samples were identified by using the R package scDblFinder (v1.6.0) with the parameter dbr.sd=1 based on the initial clustering (*67*). Cells were grouped into singlet and doublet. Cells with strong expression of hemoglobin genes (HBB, HBZ, HBM, HBQ1) were identified in individual samples by the R package scater (v1.22.0) with parameters nmad=3 and log=T. These low-quality cells and cells in clusters with a high fraction of low-quality cells were filtered out. After stringent quality control, there were 86,699 cells, and the second round of clustering was performed with the new consensus features generated using Scanorama and Seurat. Cell populations were identified based on the enriched expression of genes. For identifying subpopulations in specific cell types, additional clustering was further performed by using the functions FindNeighbors and FindClusters in the R package Seurat.

### Identification of DEGs and GO enrichment analysis

The DEGs were identified using the FindAllMarkers or FindMarkers function in Seurat. Genes with an adjusted p value less than 0.05 were considered DEGs. These DEGs with an adjusted p value less than 0.001 and a fold change of the natural logarithms greater than 0.5 were used for heatmap visualization. GO analysis was performed by using the R package clusterProfiler (v4.0.5) with the annotation R package org.Hs.eg.db (v3.13.0) (*68*).

### Identification of stage-specific gene clusters in oocytes

The R package Mfuzz (v2.52.0) was used to identify stage-specific gene clusters in oocytes (*69*). Oocytes were grouped into continuous stages according to the subpopulations identified. The average expression levels of genes for each stage were used as input for clustering. The genes were clustered into six clusters with default parameters.

### RNA velocity analysis and inferring pseudotime

RNA velocity was generated by python package velocyto with default parameters. Repeats annotation were retrived and downloaded from the UCSC genome browser. Velocity map were generated by python package scVelo, followed by inferring pseudotime with the function scv.tl.velocity_pseudotime.

### Transcriptional regulatory network analysis

The pySCENIC python package (v0.11.2) and R package hdWGCNA and WGCNA were adopted to infer the transcriptional regulatory network for a specific subpopulation in oocytes (*70*). Briefly, raw counts and a list of known TFs for humans were used as input for network inference (GRNBoost2). Cells were combined into metacells by the function construct_metacells in hdWGCNA with parameters k=25 and reduction= ‘pca’. The WGCNA analysis was performed by R package WGCNA with default parameters. The network generated by GRNBoost2 were subset by genes in cell-type specific module and cell-type specific DEGs. The network of the regulon and its targets was visualized using Cytoscape (v3.8.0) (*71*).

### Inference of cell-cell interactions based on single-cell RNA sequencing data

For inference of cell-cell interactions between oocytes and GCs and stromal cells and GCs, the CellphoneDb python package (v2.14.0) was adopted (*72*). The CellChat R package (v1.1.3) was adopted for macrophages with GCs or stromal cells (*73*). Default parameters or recommended procedures were followed.

### Cross-species integration of single-cell RNA sequencing data

For integration of single-cell RNA sequencing data from cynomolgus monkeys and humans, the FindIntegrationAnchors and IntegrateData function was adopted in the Seurat R package Seurat (*74*). Briefly, common genes were selected in our cynomolgus monkey data and two published human datasets, and raw counts for these common genes were retained. Normalization was performed in each dataset, and raw counts were scaled to 10,000. Expression levels were defined as log2 (normalized counts+1). In addition, 1,500 highly variable features were identified in each dataset, and repeated features were further selected for integration using the functions FindVariableFeatures and SelectIntegrationFeatures. The remaining procedures were followed as recommended in the Seurat documentation.

For annotation of cells from human datasets, the R package scmap was used with parameters n_feature=500 and threshold=0.5. These unassigned cells in human datasets were filtered out.

### Model for calculation of cell-type-specific aging clocks based on the monkey ovarian atlas

The development stages were regressed onto gene expression levels by using the R package glmnet (v4.1-3). For a given cell type, pseudocells were generated from different development stages and were used for LASSO regression. A total of 250 pseudocells were generated by combining raw counts for five randomly selected cells for each stage. The expression levels for pseudocells were normalized to 10,000 and log2-transformed. Genes with a nonzero regression coefficient were considered fertility-related genes. Pearson’s correlation coefficient (R) is used for measuring the strength of the relationship between predicted stages and the real developmental stages.

### H&E staining of monkey ovaries

The ovarian tissues were fixed overnight in 4% paraformaldehyde (PFA) (Sigma, P6148) at 4 °C, dehydrated with graded ethanol (70%, 80%, 95%, 100%, 100%), vitrified with dimethylbenzene, and embedded in paraffin using a tissue embedder. Paraffin blocks were sectioned (5 μm thickness) using a microtome on slides. After drying at 42 ℃ overnight, the slices were passed with a series of xylene (100% 3 times) and alcohol (100%, 100%, 95%, 80%, 70%). After brief washes in PBS, the slides were incubated with haematoxylin solution. The sections were then differentiated in 1% acid alcohol for 10 seconds and washed with PBS for 5 minutes. After incubation in eosin counterstain, the samples were subsequently dehydrated in graded ethanol (95%, 100%, 100%) and immersed in xylene 3 times for 3 minutes. The slides were then sealed with neutral resin.

### Masson’s trichrome staining of monkey ovaries

Masson’s staining was carried out according to the manufacturer’s manual (Solarbio, G1340). In brief, the paraffin sections were deparaffinized as previously described. After a brief wash in distilled water, the slides were incubated with haematoxylin iron staining solution for 10 minutes. The slides were then washed with running tap water to remove excess haematoxylin, differentiated in 1% acid alcohol for 10 seconds and washed with running tap water for 1 minute. After magenta dyeing for 10 minutes and aniline blue dyeing for 1 minute, the sections were washed with weak acid solution and dehydrated in graded ethanol (95%, 100%, 100%) for 5 seconds. Then, the sections were immersed in xylene 3 times for 1 minute and sealed with resin.

### Immunohistochemical staining of monkey ovaries

Immunohistochemical staining was carried out according to the manufacturer’s manual (ZSGB-BIO, PV-6001). In brief, the paraffin sections were deparaffinized as previously described. After PBS washes, the slides were treated for 15 minutes at 98 °C in a microwave with 0.01 M sodium citrate buffer (pH 6.0) for antigen retrieval. The sections were then washed with PBS, endogenous peroxidase blocker was added, and the sections were incubated at RT for 10 minutes. After PBS buffer rinses for 3 minutes × 3 times, the slides were incubated at 37 ℃ for 1 hour with primary antibodies and then rinsed with PBS buffer 3 minutes × 3 times. After incubation at RT for 20 minutes with secondary antibodies and washing with PBS buffer, the slides were incubated with 3,3’-diaminobenzidine (DAB) solution (ZSGB-BIO, ZLI-9018) for 5 minutes. After significant colour development, the slides were washed for 5 minutes and stained with haematoxylin solution for 5 minutes. The slides were then dehydrated and sealed as previously described.

### Immunofluorescence staining of monkey ovaries

For immunostaining, paraffin sections after rehydration were washed with PBS at RT. For antigen retrieval, sections were treated for 15 minutes at 98 °C in a microwave with 0.01 M sodium citrate buffer (pH 6.0). After cooling, the slides were rinsed three times with PBS and blocked for 1 hour at RT in blocking buffer made with 3% BSA (Merck Millipore, 0218072801) and 0.02% Triton X-100 (Sigma, 9002-93-1) dissolved in PBS. Subsequently, sections were incubated at 4 ℃ overnight with primary antibodies, followed by 1 hour with secondary antibodies at RT. All samples were diluted in blocking buffer. Finally, the slides were sealed with anti-fluorescence quenching agent and stored at −20 ℃ for further imaging.

### Examination of androgen levels in peripheral blood

Serum samples were also collected from 36 other cynomolgus monkeys aged 1–27 years. Enzyme-linked immunosorbent assays (ELISAs) was used to quantify androgen (Beijing Zhichao Weiye Biological Technology, ZC70048) levels in serum. The detection system consisted of a biotinylated secondary antibody and horseradish peroxidase labelled with streptavidin. Specimens were assayed in triplicate.

## Acknowledgements

We thank S.W. L., X. L. Z. and the imaging platform for their outstanding support. This work was supported by the National Key Research and Development Program of China (2019YFA0110900, 2022YFA1104100, 2018YFA0107701), the Strategic Priority Research Program of the Chinese Academy of Sciences (XDA16021400, XDA16020700), the CAS Project for Young Scientists in Basic Research (YSBR-012) and the Informatization Plan of Chinese Academy of Sciences (CAS-WX2021SF-0101).

## Author contributions

F. Guo, H. W., F. Gao and L. Y. designed the study and supervised the overall experiments. Y. Z., F. Z., W. T. and Q. Y. collected all of the ovarian samples. L. Y., Y. Z., X. L., J. Q. and J. Z. isolated ovarian single cells and established the scRNA-seq library. L. Y., Y. Z., X. L., Y. H. and B. H. executed the original data management and quality control. X. L., Y. Z., M. W. and X. J. performed the dataset analysis. F. Guo, H. W., F. Gao, L. Y., Y. Z., X. L. and Y. S. wrote the manuscript.

## Supplementary materials

Fig. S1-12

## Data and materials availability

The sequencing data generated in this study will be made publicly available upon publication.

## Supplementary Materials for

**Fig. S1.**
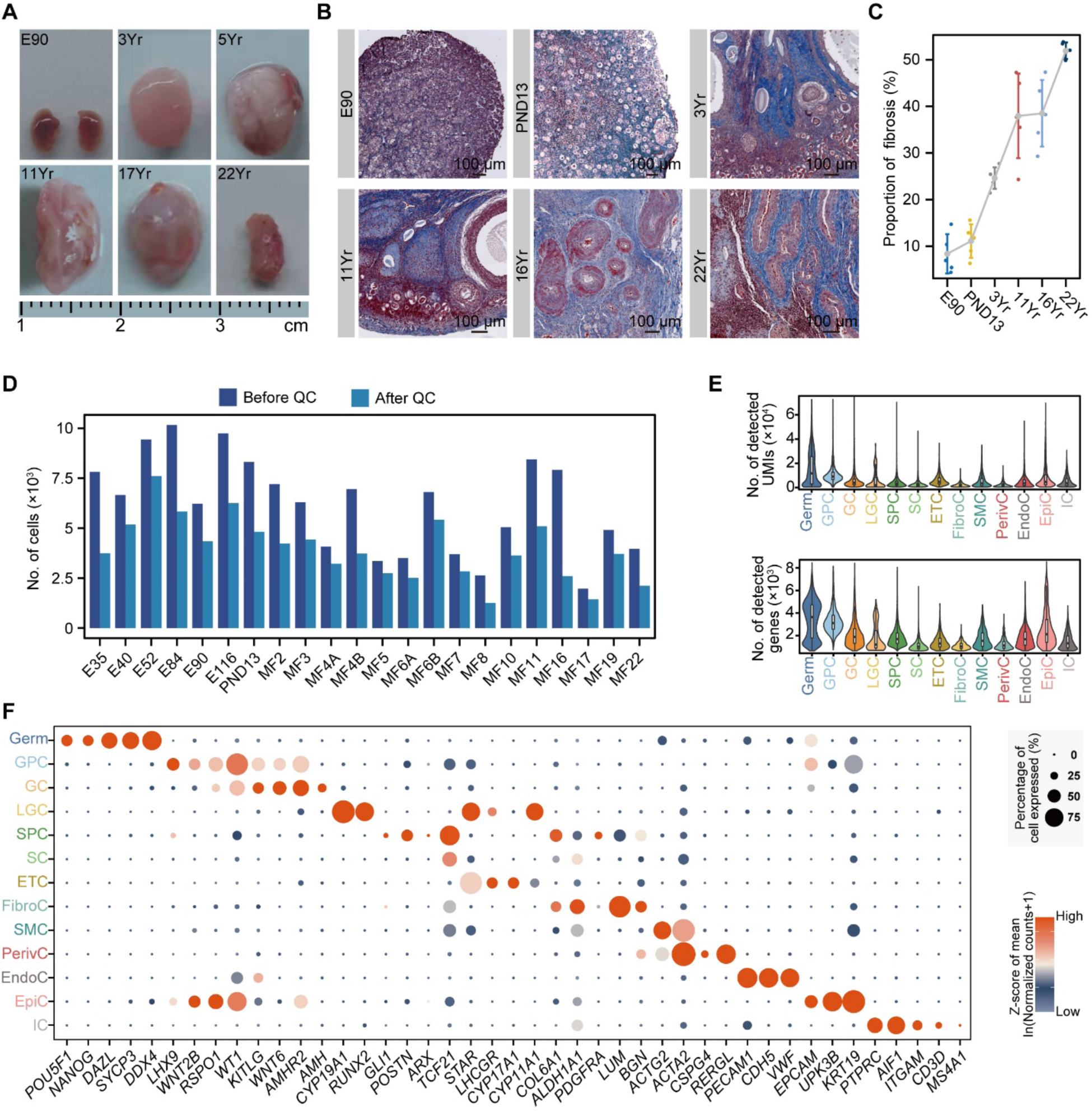
Ovarian signatures of cynomolgus monkeys and gene markers for each ovarian cell cluster. **A,** Representative images of sampled ovaries. Scale division is as indicated. **B,** Masson trichrome staining of ovaries of different ages. Scale bars are as indicated. **C,** Proportions of fibrosis in ovarian sections at different ages. n=5 sections. In the error bar plots, mean±SE was used for visualizing errors. **D,** Bar plot showing the number of cells sequenced and used for the individual sample in this study. **E,** Violin plot showing the number of detected UMIs (left) and genes (right) in an individual cell. Germ, GPC, GC, LGC, SPC, SC, ETC, FibroC, SMC, PerivC, EndoC, EpiC and IC denote germ cells, granulosa progenitor cells, granulosa cells, luteal granulosa cells, stromal progenitor cells, stromal cells, endocrine theca cells, fibroblasts, smooth muscle cells, perivascular cells, endothelial cells, epithelial cells and immune cells, respectively. For boxplot in violin plots, the lower boundaries of the box indicate the 25th percentiles of expression level and the upper boundaries indicate the 75th percentiles. The line inside the box indicates the median. The ends of the whiskers indicate the highest data values within 1.5 × IQR of the 75th percentile value and the lowest data values within 1.5 × IQR of the 25th percentile value. **F,** Dot plot depicting the relative mean expression levels of representative genes identified in the indicated cell populations in Fig. 1C. Mean expression levels are derived from logarithm-scaled normalized counts based on the single-cell RNA-seq data. The percentage of cells that express the respective gene is indicated by the size of the dot. Germ, GPC, GC, LGC, SPC, SC, ETC, FibroC, SMC, PeriC, EndoC, EpiC and IC denote germ cells, granulosa progenitor cells, granulosa cells, luteal granulosa cells, stromal progenitor cells, stromal cells, endocrine theca cells, fibroblasts, smooth muscle cells, perivascular cells, endothelial cells, epithelial cells and immune cells, respectively.

**Fig. S2.**
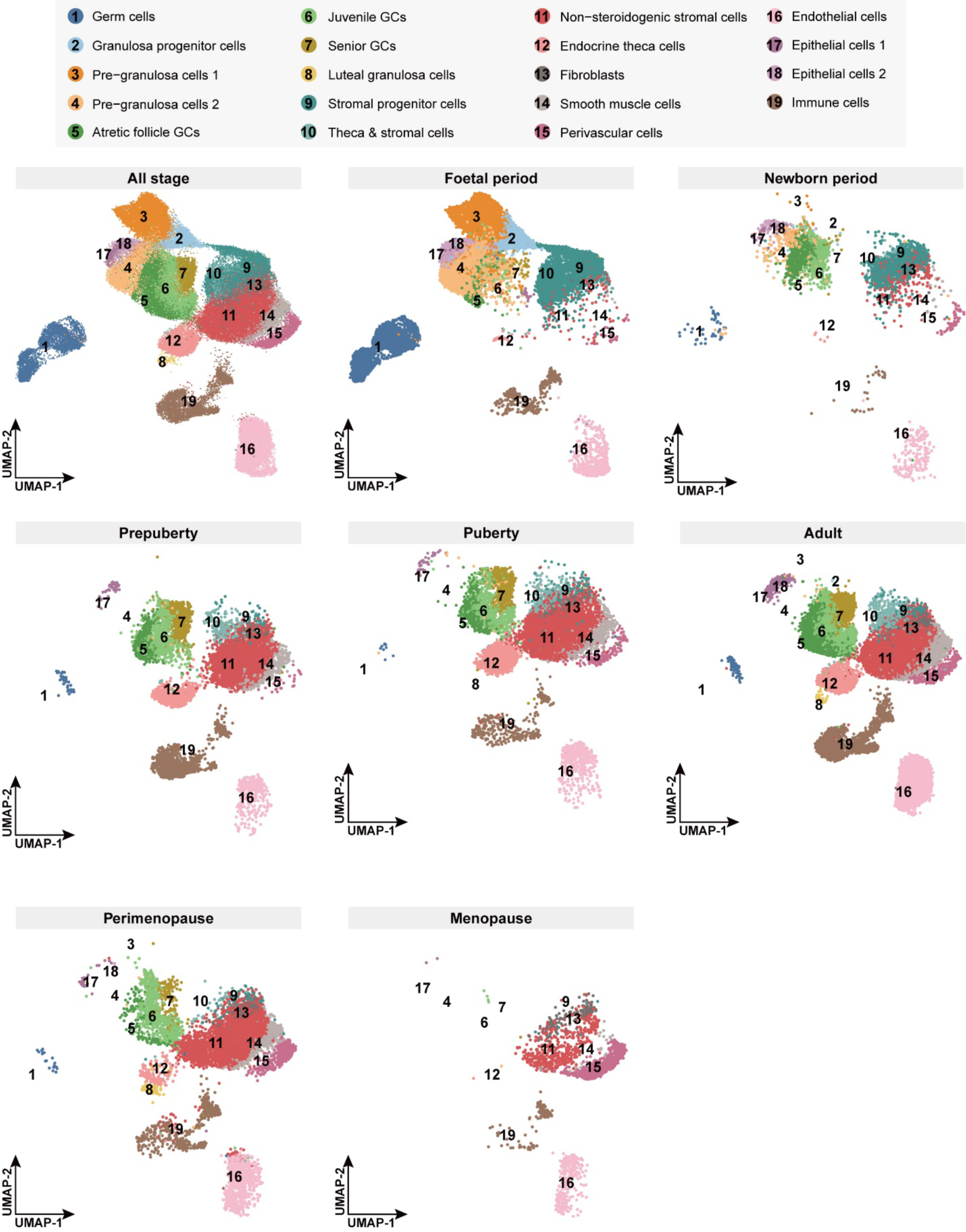
Cellular composition of ovarian cells across the foetal, newborn, prepuberty, puberty, adult, perimenopause, and menopause stages. UMAP embedding visualization for 19 cell clusters identified in ovaries of cynomolgus monkeys in each stage.

**Fig. S3.**
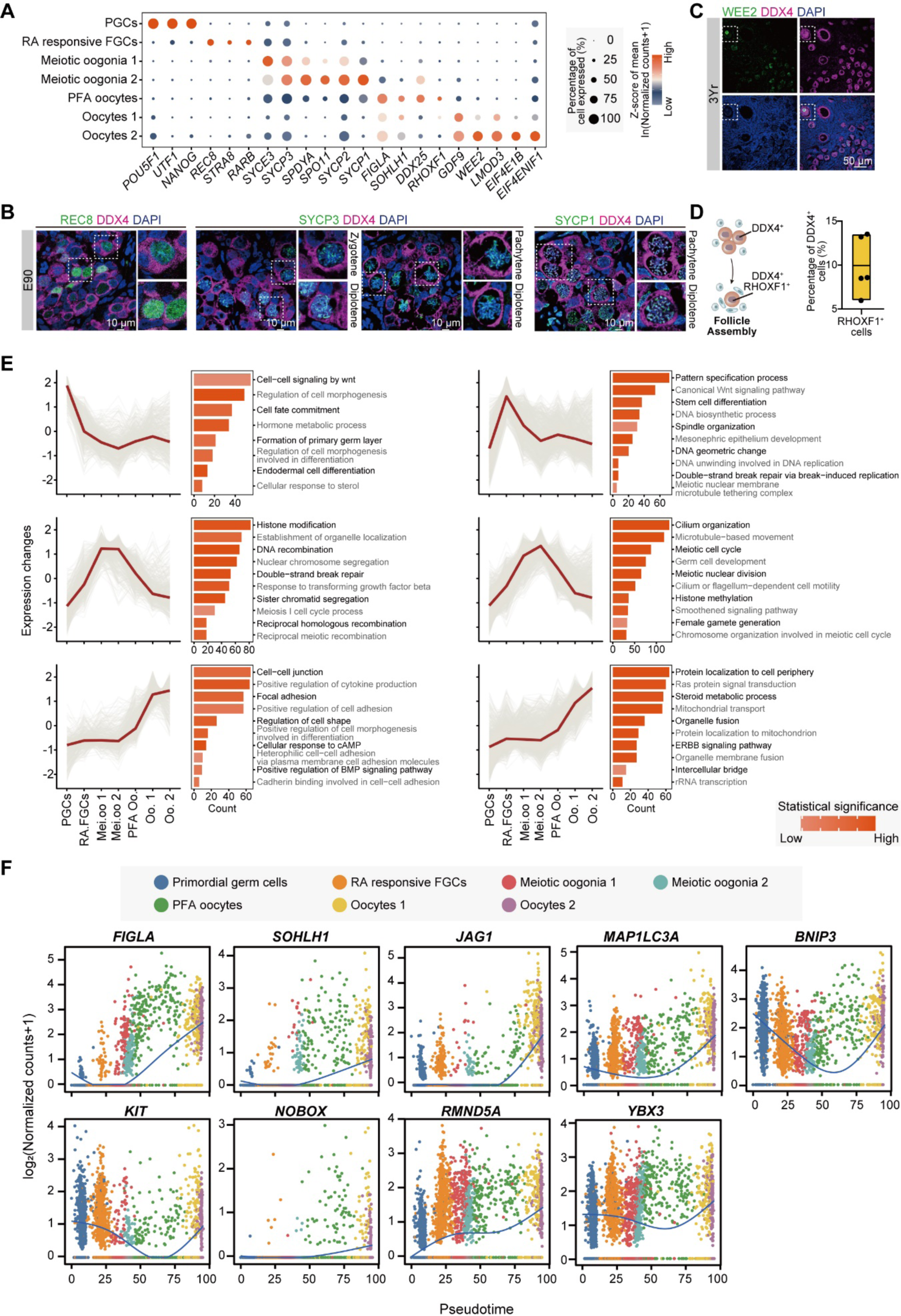
Validation of oocyte subtypes and identification of their transcriptomic characteristics. **A,** Dot plot depicting the relative average expression levels of representative genes identified in the indicated cell populations in germ cells. Average expression levels were derived from logarithm-scaled normalized counts based on the single-cell RNA-seq data. The percentage of cells that express the respective gene is indicated by the size of the dot. PGCs denotes primordial germ cells and FGCs for foetal germ cells, RA for retinoic acid and PF primordial follicle. **B,** Immunofluorescence staining for REC8 (marker of RA-responsive FGCs), SYCP3 (marker of meiotic oogonia 1), SYCP1 (marker of meiotic oogonia 2) and DDX4 in ovarian tissues from E90 cynomolgus monkeys. The nuclei were counterstained with DAPI. RA-responsive FGC and meiotic oocytes in various stages (framed by dashed lines) are shown, and magnified images are shown on the right. RA denotes retinoic acid; FGC denotes foetal germ cells, and PF denotes primordial follicles. Scale bars are as indicated. **C,** Immunofluorescent images of ovarian tissue from 3 Yr cynomolgus monkeys stained with anti-WEE2 (marker of oocytes in growing follicles) and anti-DDX4 antibodies. The nuclei were counterstained with DAPI. The expression of WEE2 is detected as a marker of growing oocytes (framed by dashed). Scale bars are as indicated. **D,** Proportion of PF assembly oocytes indicated by DDX4^+^ and RHOXF1^+^ in the ovaries of E90 cynomolgus monkeys. For boxplot plots, the lower boundaries of the box indicate the 25th percentiles of expression level and the upper boundaries indicate the 75th percentiles. The line inside the box indicates the median. **E,** Representative gene cluster identified in subpopulations of germ cells. Genes were clustered by their expression pattern along the developmental stage of oocytes by using the R package Mfuzz. Representative GO terms for these genes are shown on the right. RA denotes retinoic acid; Mei. Oo denotes meiotic oogonia; PFA Oo denotes primordial follicle assembly oocytes. **F,** Trend lines showing relative expression levels for representative genes in subpopulations identified in oocytes. RA denotes retinoic acid; FGC denotes foetal germ cells; PFA denotes primordial follicle assembly.

**Fig. S4.**
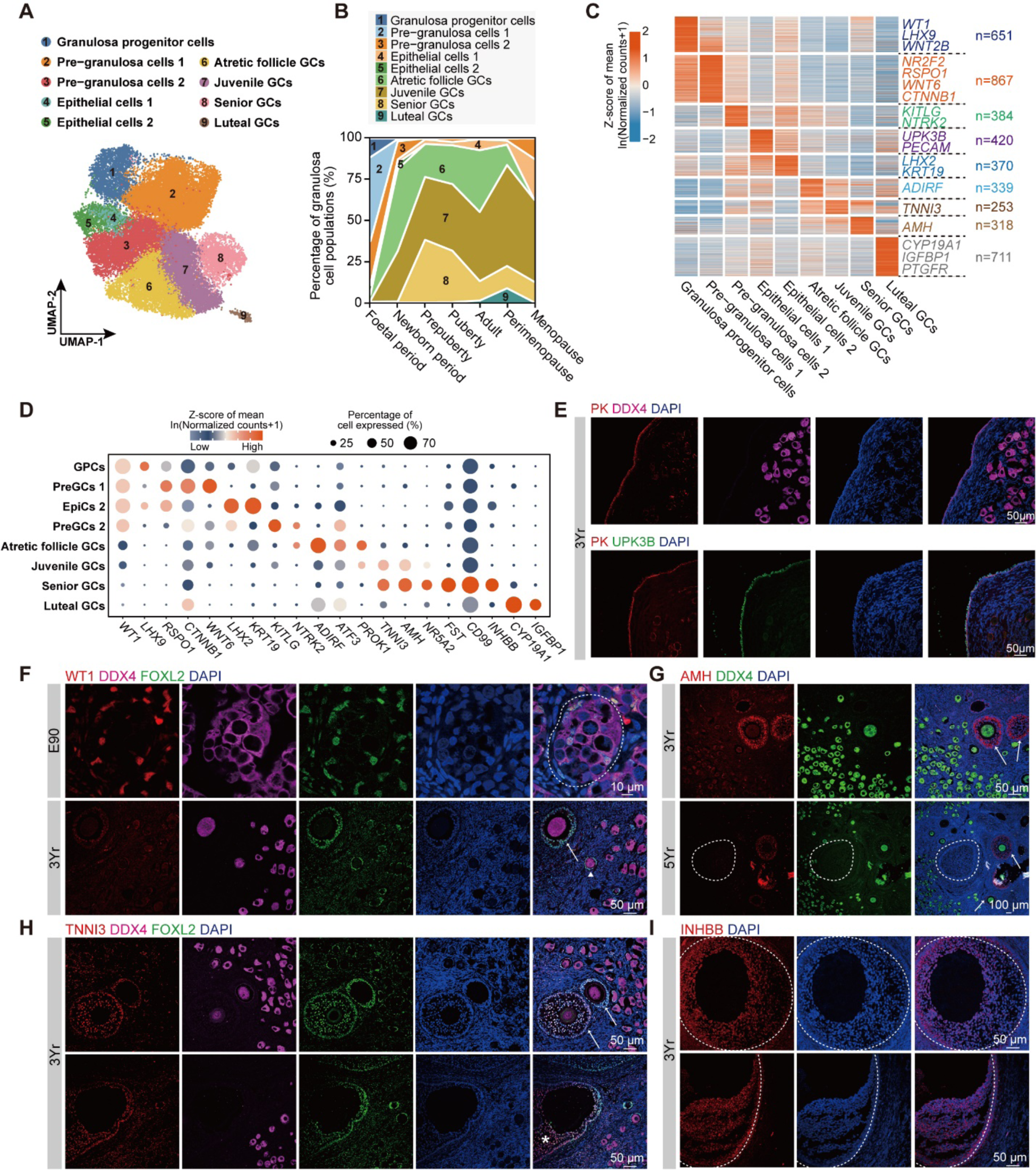
Subtypes of GCs and gene expression signatures. **A,** UMAP embedding visualization for nine subpopulations in granulosa cells and epithelial cells identified in Fig. 1C. Cells are colour coded by the subpopulations identified (left) and developmental stage for cynomolgus monkeys where cells were from (right). **B,** Stacked area plot showing the fraction of subpopulations in ovarian GCs in cynomolgus monkeys during the developmental stages and aging. GC denotes granulosa cells. **C,** Heatmap showing representative DEGs identified in different subpopulations. Gene expression levels are averaged and scaled. Cell populations are colour coded as (**A**). GC denotes granulosa cells. **D,** Dot plot depicting the relative mean expression levels of representative genes in the indicated cell subpopulations in GCs. Mean expression levels are derived from logarithm-scaled normalized counts based on the single-cell RNA-seq data. The percentage of cells that express the respective gene is indicated by the size of the dot. GC denotes granulosa cells. **E,** Immunofluorescent images of the ovary of a 3-year-old cynomolgus monkey stained with anti-Pan-Keratin (PK, marker of epithelial cells), anti-DDX4 and anti-UPK3B (marker of epithelial cells). The nuclei were counterstained with DAPI. Scale bars are as indicated. **F,** Immunofluorescence staining for WT1, DDX4 and FOXL2 in ovarian tissues from cynomolgus monkeys. The nuclei were counterstained with DAPI. The dashed circle indicates the germ cell cyst, white arrowheads indicate primary follicles, and white arrows indicate secondary follicles. The age of individual monkeys and the scale bars are as indicated. **G,** Immunofluorescence staining for AMH and DDX4 in ovarian tissues from cynomolgus monkeys. The nuclei were counterstained with DAPI. The white arrows point to secondary follicles, and the dashed line marks the boundary of the atretic follicle. The age of individual monkeys and the scale bars are as indicated. **H,** Immunofluorescence staining for TNNI3, DDX4 and FOXL2 in ovarian tissues from cynomolgus monkeys. The nuclei were counterstained with DAPI. White arrows point to secondary follicles, and asterisks indicate antral follicles. The age of individual monkeys and the scale bars are as indicated. **I,** Immunofluorescence staining for INHBB (marker of GCs in antral follicles) in 3 Yr and 5 Yr ovarian tissues from cynomolgus monkeys. The nuclei were counterstained with DAPI. The dashed line marks the boundary of the antral follicles. Scale bars are as indicated.

**Fig. S5.**
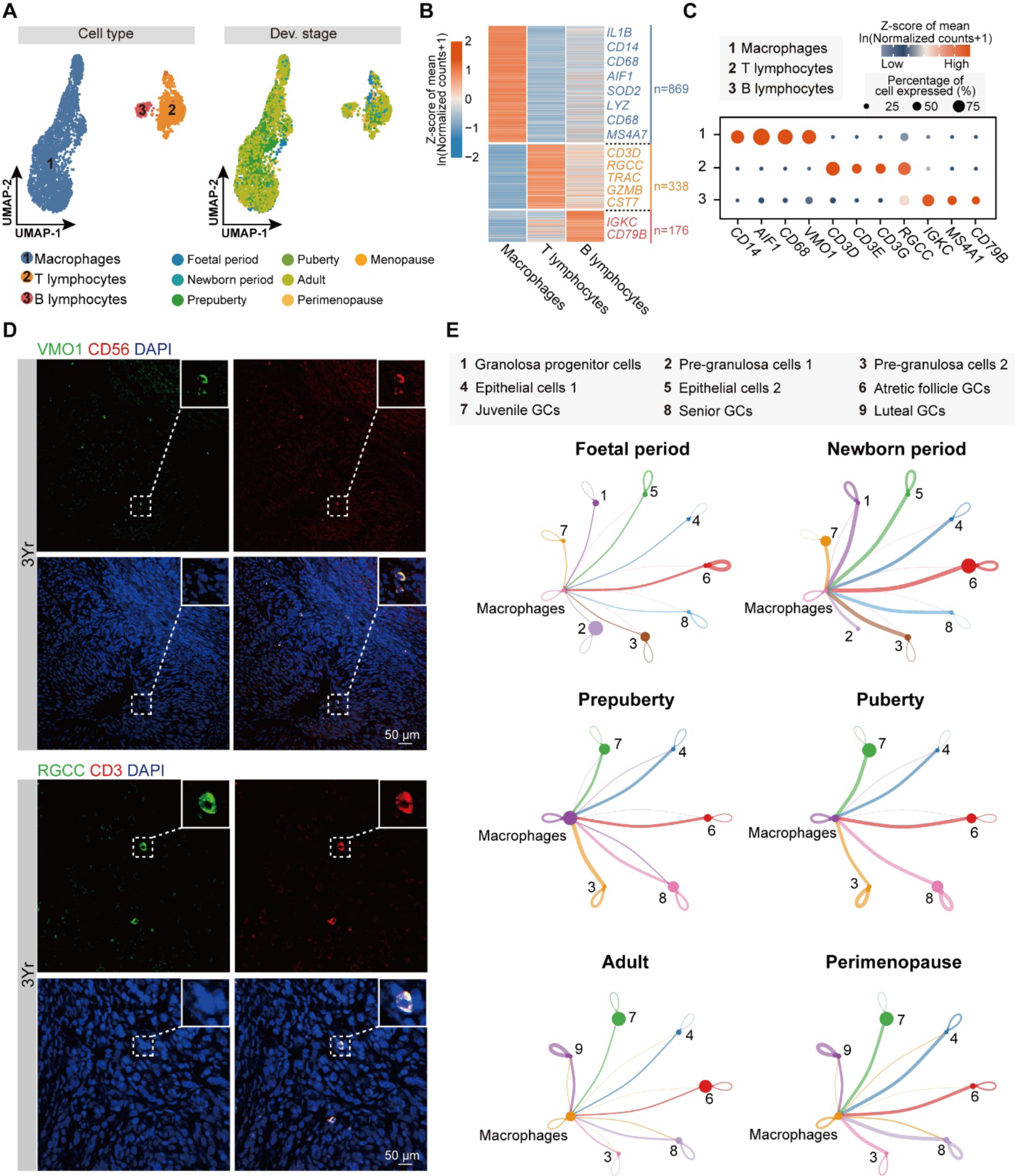
Immune cell populations and their interactions with GCs. **A,** UMAP embedding visualization for subpopulations in immune cells identified in Fig. 1C. Cells are colour coded by subpopulations identified (left) and developmental stage for cynomolgus monkeys where cells were from (right). **B,** Heatmap showing representative DEGs identified in macrophages, T lymphocytes and B lymphocytes. Gene expression levels are averaged and scaled. **C,** Dot plot depicting the mean expression levels of representative genes identified in macrophages, T lymphocytes, and B lymphocytes. Mean expression levels are derived from logarithm-scaled normalized counts based on the single-cell RNA-seq data. The percentage of cells that express the respective gene is indicated by the size of the dot. **D,** Immunofluorescence staining for VMO1 (marker of macrophages), CD56 (marker of macrophages), RGCC (marker of T lymphocytes) and CD3 (marker of T lymphocytes) in ovarian tissues of the 3 Yr cynomolgus monkey. The nuclei were counterstained with DAPI. Macrophages and T lymphatic cells are indicated (framed by dashed lines), and magnified images are shown in the top right corner. Scale bars are as indicated. **E,** Circle plots showing interactions between GC subpopulations and macrophages. These interactions were inferred by using the R package CellChat. The width of edges represents connectivity between clusters, and the size of nodes represents the number of cells in the indicated cluster.

**Fig. S6.**
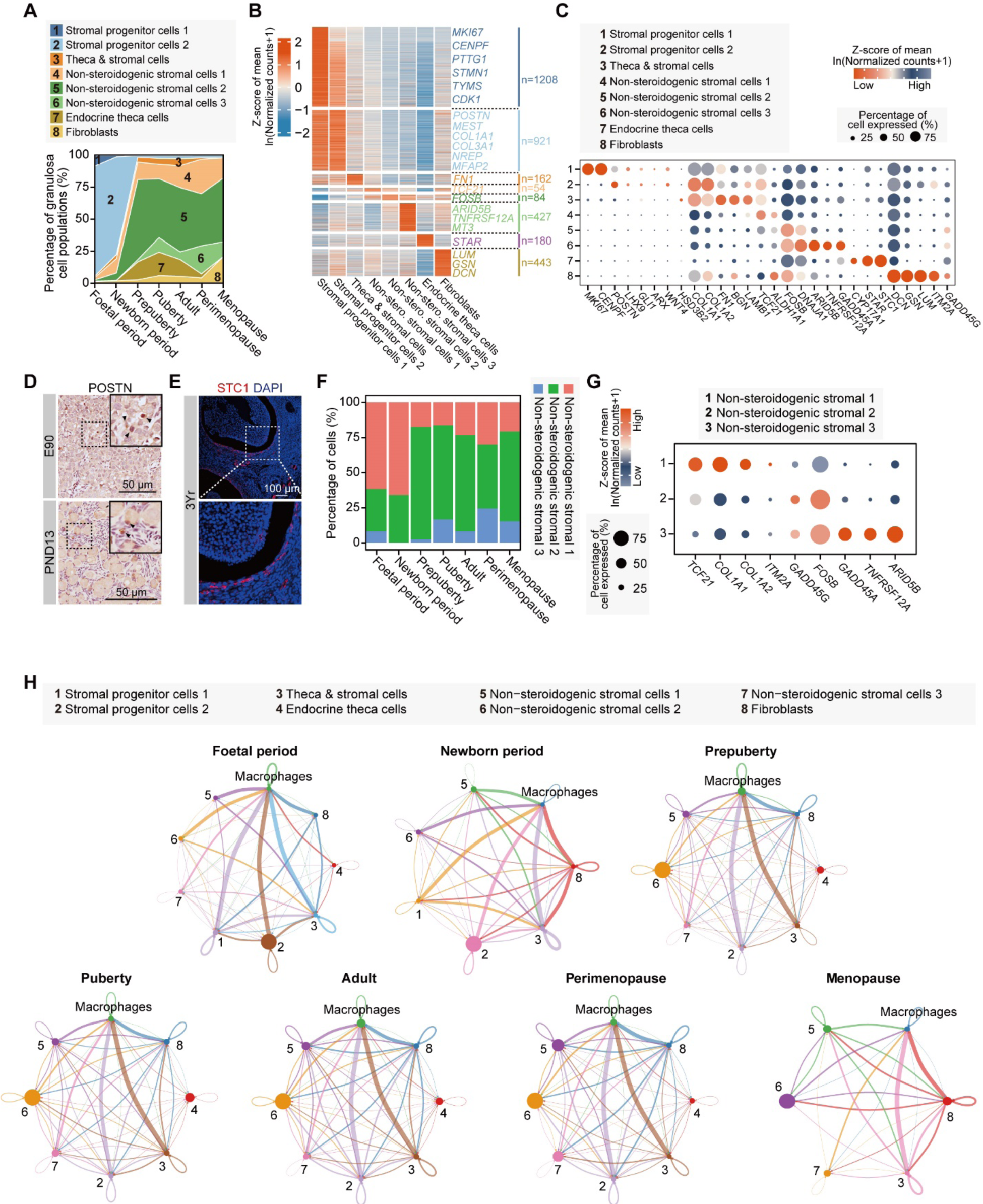
Characterization of transcriptomic features of stromal cells and their interaction with immune cells. **A,** Stacked area plot showing the fraction of cell subpopulations identified in stromal cells in cynomolgus monkeys during developmental stages and aging. **B,** Heatmap showing representative DEGs identified in different subpopulations. Gene expression levels are averaged and scaled. Cell populations are colour coded as (**A**). Nonstero. denotes nonsteroidogenic. **C,** Dot plot depicting the relative average expression levels of representative genes identified in eight subpopulations. Average expression levels are derived from logarithm-scaled normalized counts based on the single-cell RNA-seq data. The percentage of cells that express the respective gene is indicated by the size of the dot. **D,** Immunohistochemistry results for POSTN (marker of stromal progenitor cells) in ovarian tissues from cynomolgus monkeys. The dashed frames indicate SPCs, and magnified images are at the top right corner. The age of individual monkeys and the scale bars are as indicated. **E,** Immunofluorescent images of the ovary of 3 Yr cynomolgus monkeys stained with anti-STC1 (marker of endocrine theca cells, hereinafter inclusive). The nuclei were counterstained with DAPI. The expression of STC1 was detected as a marker of endocrine theca cells (framed by dashed), magnified image on the right. Scale bars are as indicated. **F,** Bar plot showing the percentage of subpopulations in nonsteroidogenic cells in different developmental stages. **G,** Dot plot depicting the mean expression levels of representative genes identified in subpopulations in nonsteroidogenic cells. Mean expression levels are derived from logarithm-scaled normalized counts based on the single-cell RNA-seq data. The percentage of cells that express the respective gene is indicated by the size of the dot. **H,** Circle plots showing the interaction between theca and stromal cell subpopulations and macrophages. These interactions were inferred by using the R package CellChat. The width of edges represents connectivity between clusters, and the size of nodes represents the number of cells in the indicated cluster. ECM denotes the extracellular matrix.

**Fig. S7.**
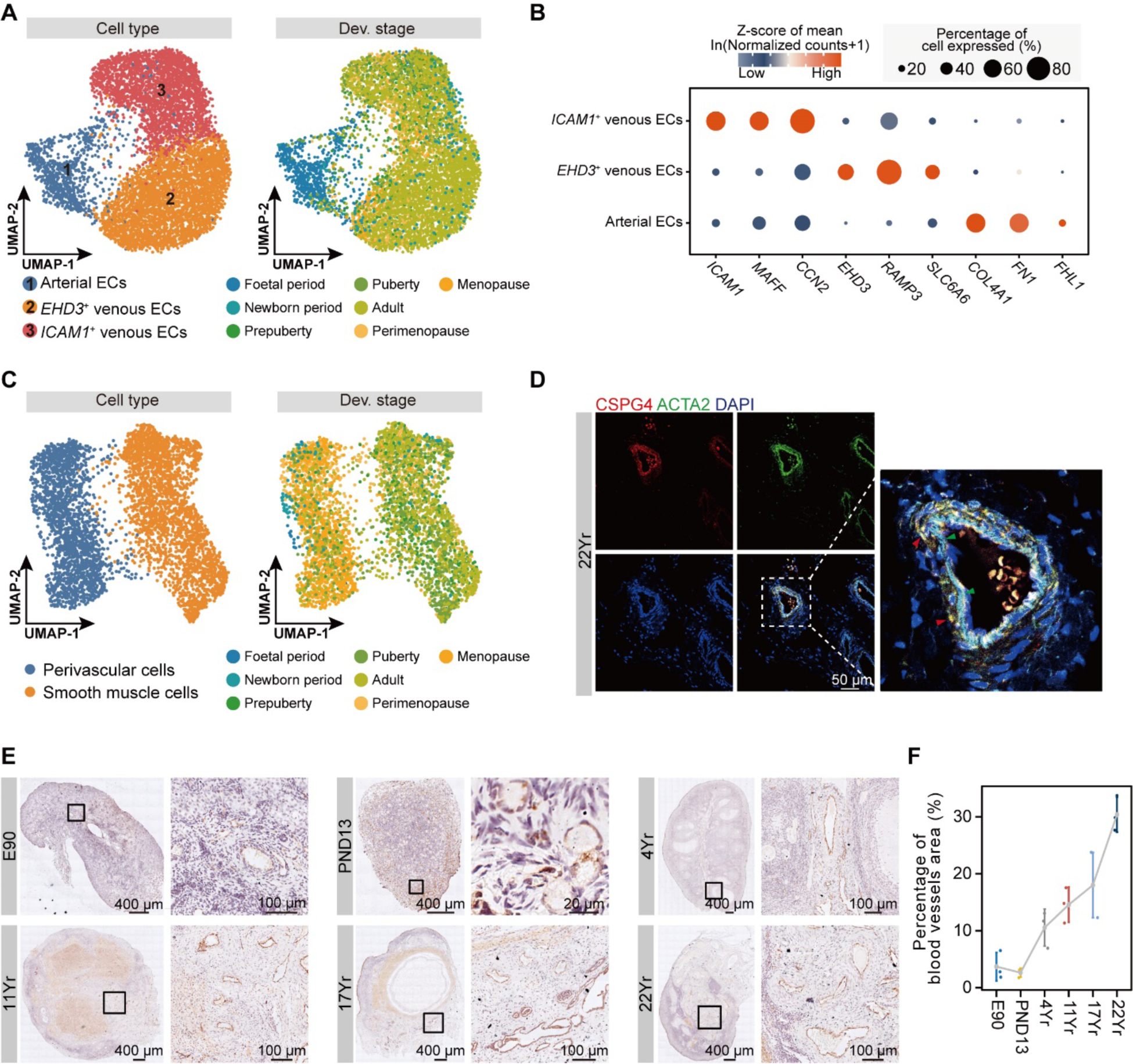
Examination of epithelial, endothelial, smooth muscle and perivascular cells in ovaries of cynomolgus monkeys. **A,** UMAP embedding visualization for three subpopulations in endothelial cells identified in Fig. 1C. Cells are colour coded by subpopulations identified (left) and developmental stage for cynomolgus monkeys where cells are from (right). EC denotes endothelial cells. **B,** Dot plot depicting the relative mean expression levels of representative genes identified in subpopulations in endothelial cells. Mean expression levels are derived from logarithm-scaled normalized counts based on the single-cell RNA-seq data. The percentage of cells that express a gene is indicated by the size of the dot. EC denotes endothelial cells. **C,** UMAP embedding visualization for smooth muscle cells and perivascular cells identified in Fig. 1C. Cells are colour coded by cell type (left) and developmental stage for cynomolgus monkeys where cells are from (right). **D,** Immunofluorescence staining for CSPG4 (a marker of perivascular cells) and ACTA2 (a marker of perivascular and smooth muscle cells) in ovarian tissues from cynomolgus monkeys. The nuclei were counterstained with DAPI. The age of individual monkeys and the scale bars are as indicated. Red triangles indicate CSPG4^+^ perivascular cells, green triangles indicate CSPG4^-^ smooth muscle cells **E,** Immunohistochemistry staining for CD31 (marker of endothelial cells) in ovarian tissues from cynomolgus monkeys. Magnified images of local blood vessels are shown on the right. The age of individual monkeys and the scale bars are as indicated. **F,** Variation in blood vessel density in ovaries from cynomolgus monkeys at different age stages. In the error bar plots, mean±SE was used for visualizing errors.

**Fig. S8.**
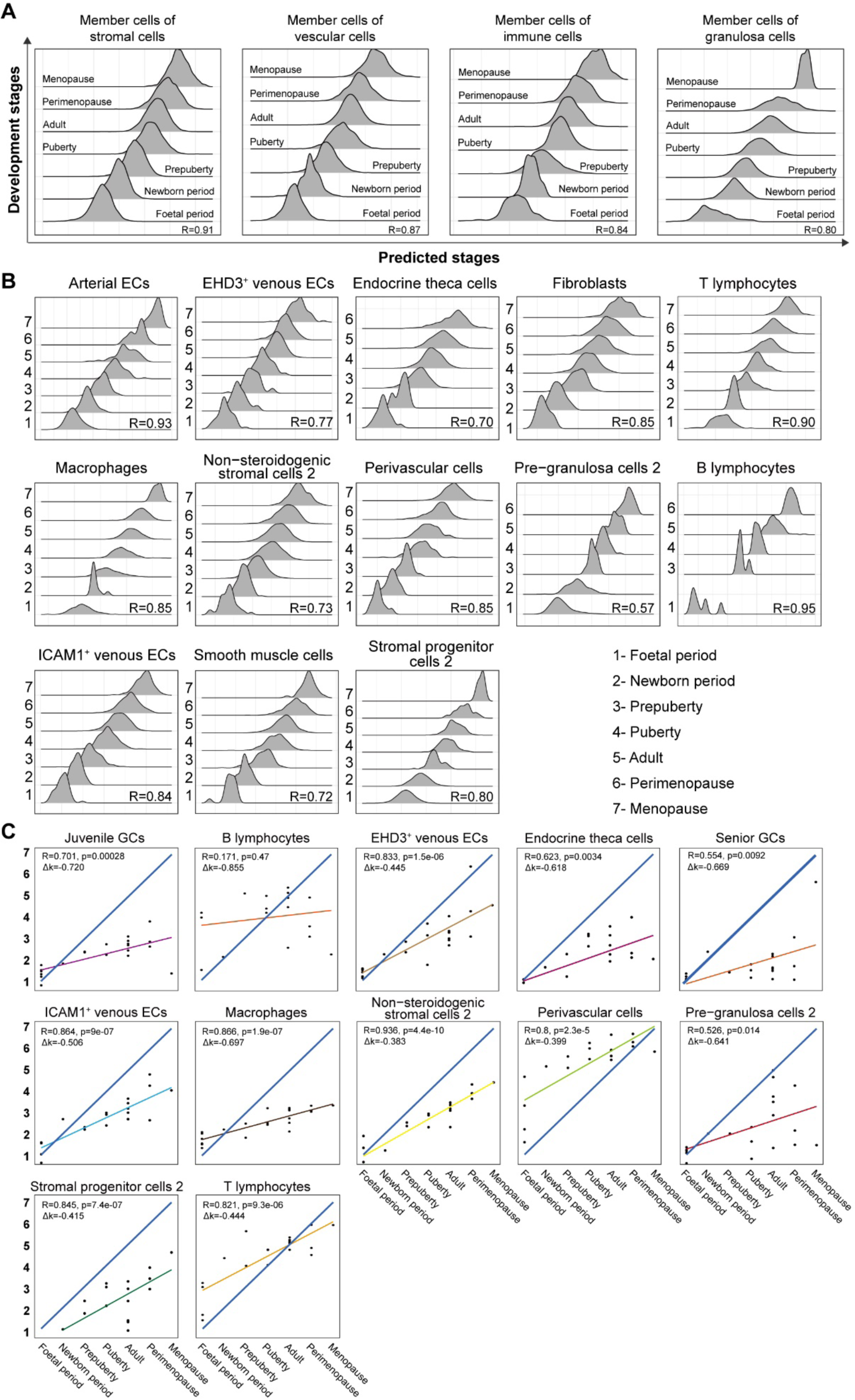
Predication of cell-type-specific developmental stages. **A,** Ridge plots showing associations between developmental stages and predicted stages based on the indicated aging clocks in stromal cells, vascular cells, immune cells and granulosa cells. **B,** Ridge plots showing associations between developmental stages and predicted stages based on the indicated aging clocks in individual cell types. **C,** Scatter plot showing correlations for average expression levels of whole ovary and a specific cell type. The R value and delta k are as indicated.

**Fig. S9.**
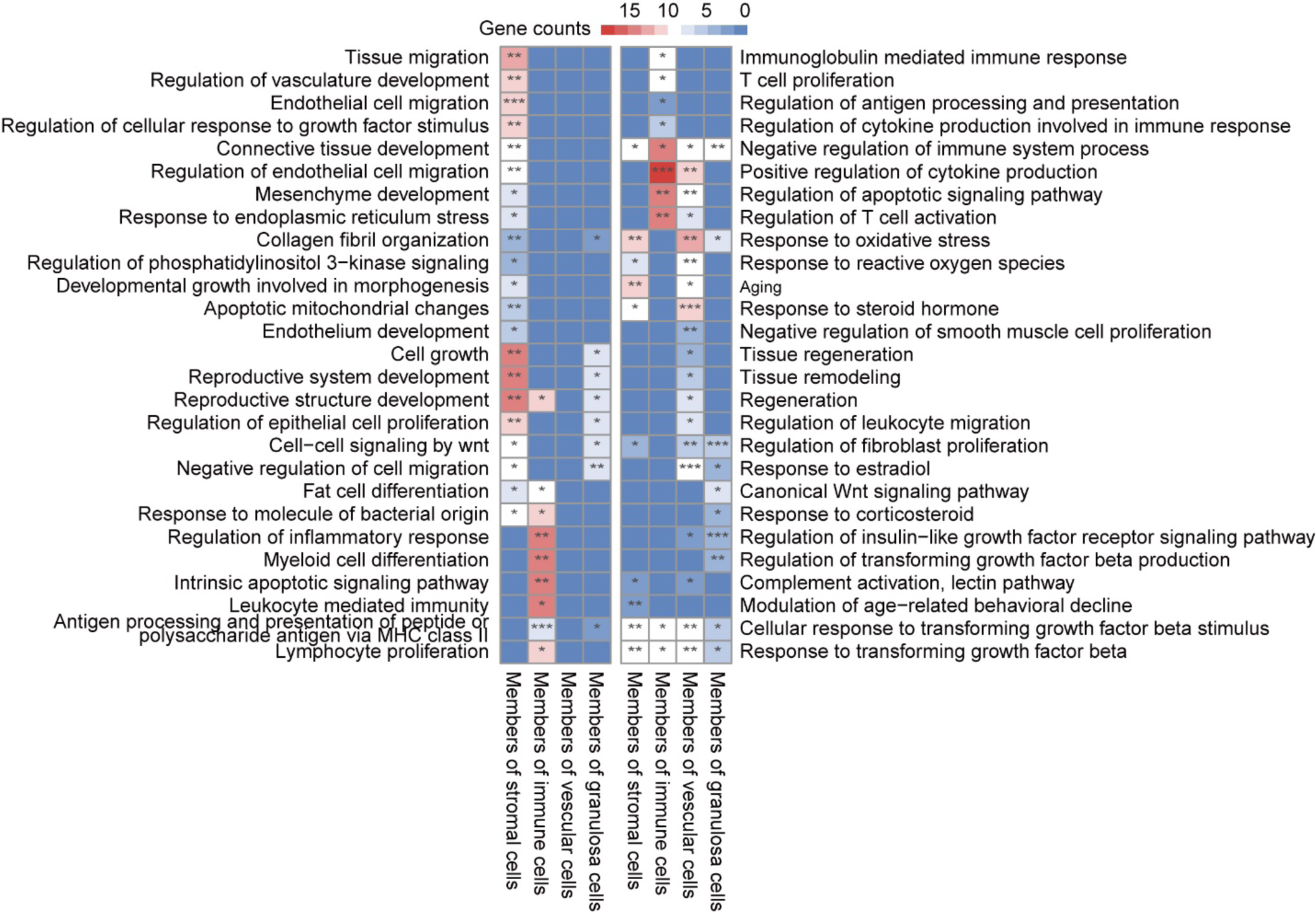
Gene ontology enrichment analysis for genes in aging clocks. Heatmap showing representative GO terms for genes used in the aging clock (stromal cells, vascular cells, immune cells and granulosa cells).

**Fig. S10.**
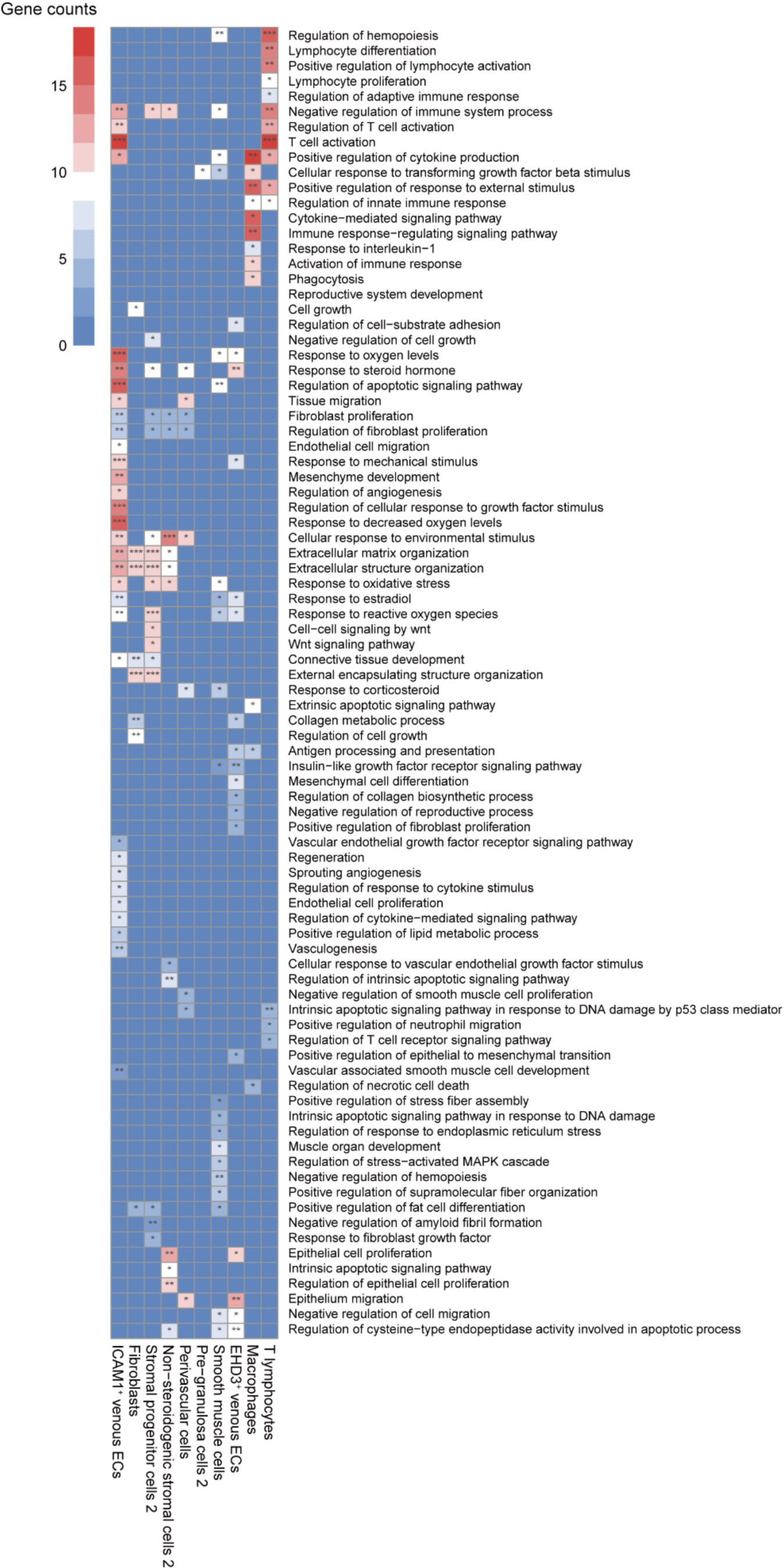
Gene ontology enrichment analysis for genes in aging clocks. Heatmap showing representative GO terms for genes used in the aging clock (individual cell types in Fig. S8**B**).

**Fig. S11.**
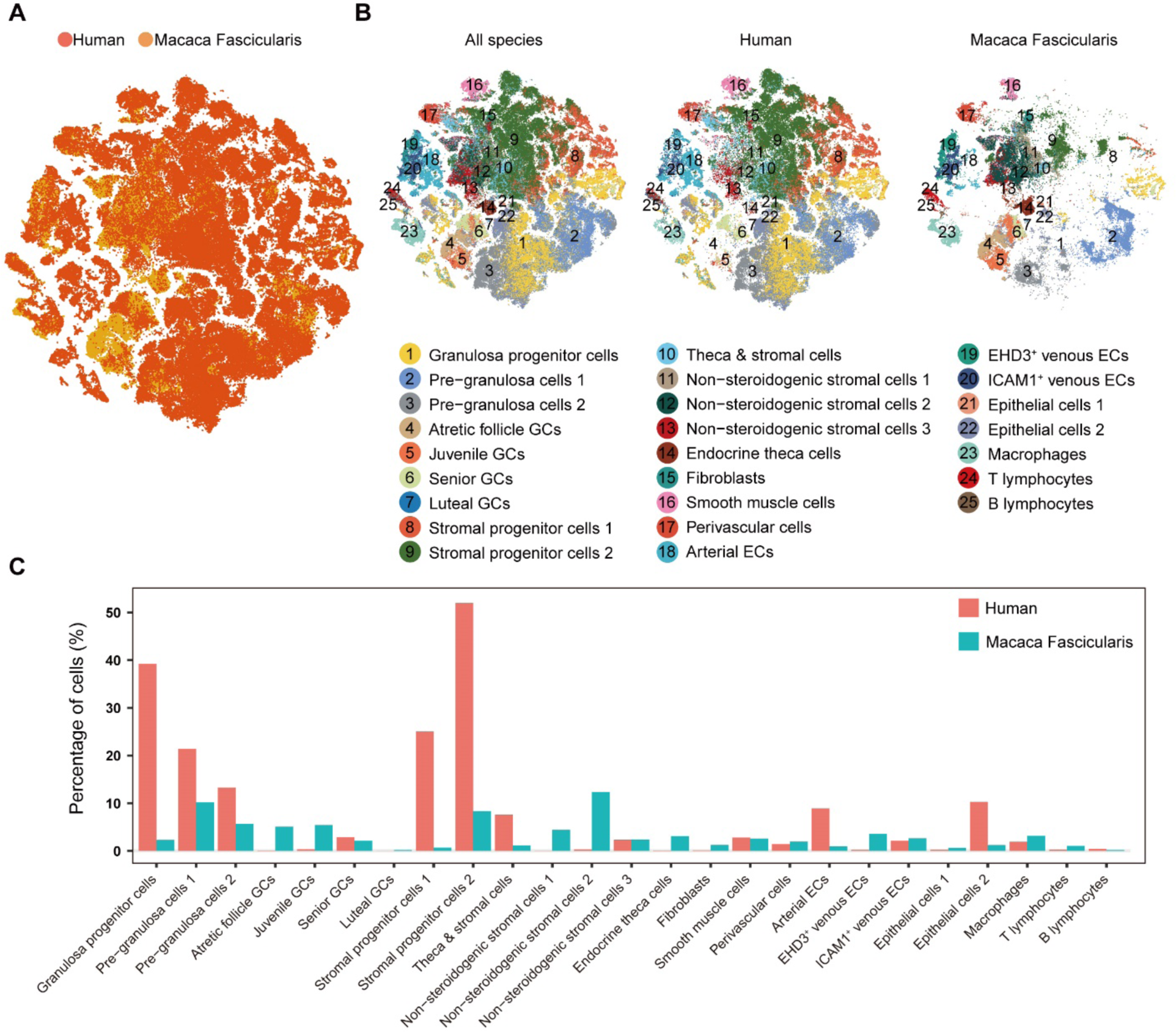
Interspecies integrated analysis of the single-cell transcriptome of cynomolgus monkey and human ovaries. **A,** t-distributed stochastic neighbour embedding (t-SNE) visualization for integrated monkey and human single-cell ovarian transcriptome data (*26, 73*). Cells are coloured by species. **B,** t-SNE visualization for integrated monkey and human single-cell ovarian transcriptome data (*26, 73*). Cells are coloured according to the different subpopulations of cells identified in this study. GC and EC denote granulosa cells and endothelial cells, respectively. **C,** Grouped bar plot showing the percentage of the indicated subpopulation in monkeys and humans based on integrated cross-species analysis. GC and EC denote granulosa cells and endothelial cells, respectively.

**Fig. S12.**
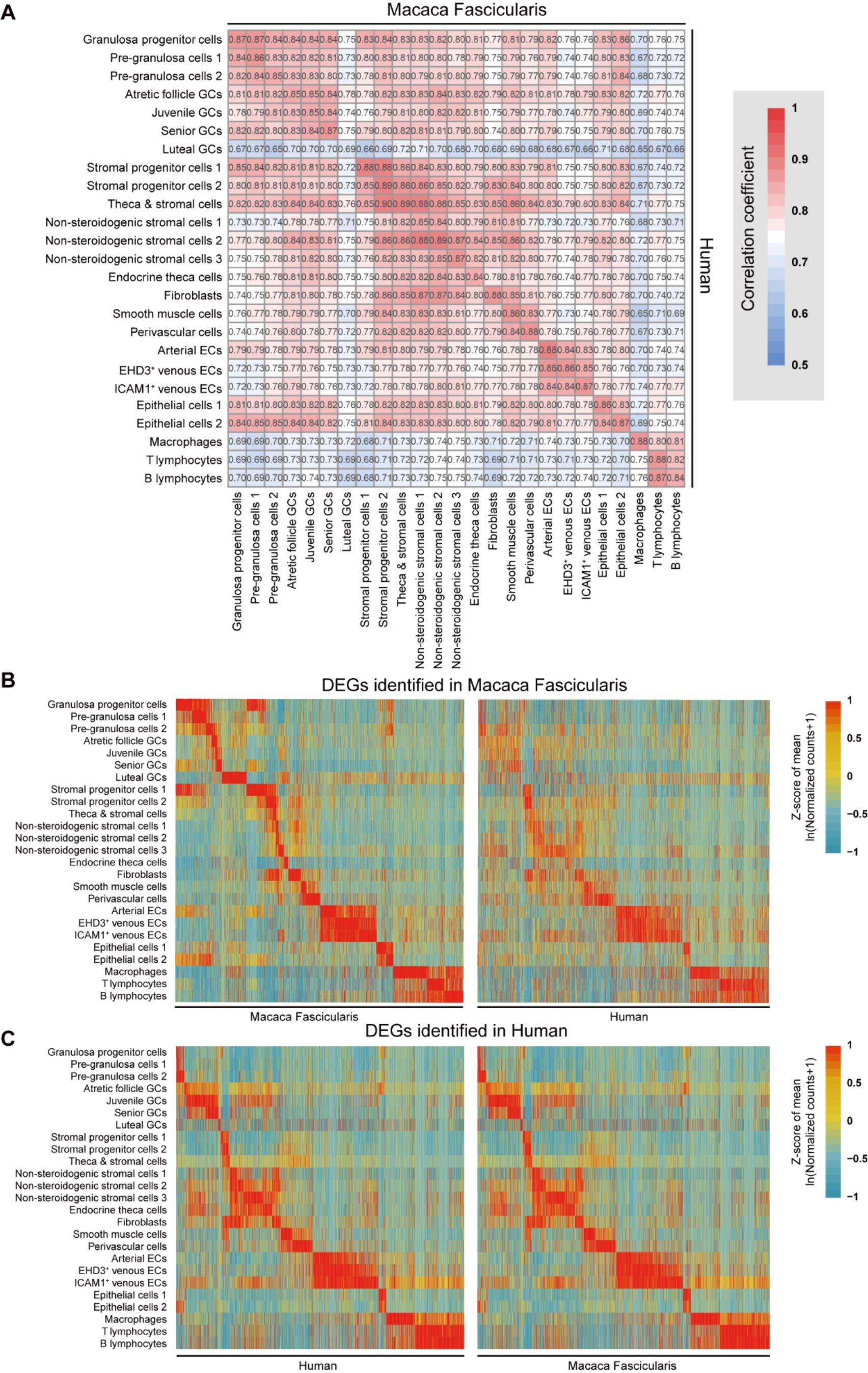
Comparison of cellular diversity and gene expression of ovarian cells between monkeys and humans. **A,** Heatmap showing Pearson correlation coefficients among cell subpopulations identified in this study. GC and EC denote granulosa cells and endothelial cells, respectively. **B,** Heatmap showing the expression levels in cynomolgus monkeys and humans for DEGs identified in the indicated subpopulation in cynomolgus monkeys. **C,** Heatmap showing expression levels in cynomolgus monkeys and humans for DEGs identified in the indicated subpopulation in humans.

